# Autobehaver: An AI-Based Pipeline for Animal Behavior Analysis

**DOI:** 10.64898/2026.05.12.724596

**Authors:** R.S. O’Neill, S. Aviles, N.M. Rusan

## Abstract

Behavior arises from the complex interplay between an organism’s nervous system, its genetic makeup, and the environment. High-resolution, high-throughput behavioral quantification is essential for dissecting biological function and the effects of genetic perturbation, but automated analysis remains challenging. Here, we present Autobehaver, an automated behavioral analysis pipeline based on a low-cost, high-throughput recording platform that captures videos of individual *Drosophila*. From each video, we extracted keypoints and used a custom Transformer to assign frame-wise behavior and orientation labels. We then converted these predictions into high-dimensional per-animal feature vectors and trained XGBoost ensembles to classify animals and identify the features that separated groups. By applying SHAP analysis to the classifier ensemble, we identified the behavioral features most informative for distinguishing groups of flies. We demonstrated the approach in several ways. First, we recovered known behavioral changes associated with heat-activated dTrpA1 activity in specific neural circuits. Second, we detected age-associated behavioral changes consistent with gradual impairment of locomotor and climbing ability. Finally, we used Autobehaver’s classifier ensemble to place animals with intermediate phenotypes along a behavioral axis and used feature-importance analysis to reveal the behavioral features underlying those intermediate states. Together, Autobehaver provides an interpretable framework for quantitative behavioral phenotyping and comparative analysis of complex genotypes.

## Introduction

*Drosophila melanogaster* has long been a powerful model organism for understanding factors that influence behavior^1^. Neuroscientists have used flies to investigate how specific neurons and complex neural circuits control behavior^2–6^. Similarly, fly models of human diseases often recapitulate behavioral traits observed in patients^7–11^.

Researchers use a variety of assays and recording platforms to study behavior in *Drosophila*. Many assays assess specific behaviors, such as climbing, locomotion, geotaxis, phototaxis, seizure sensitivity, and mating, often using a stimulus-response paradigm^12^. Other approaches rely on recording flies, sometimes in a semi-high-throughput manner, and analyzing the recordings for specific behaviors including grooming^13^, social behavior^14^, aggression^15^, flight take-off^16^, gait^17^, and sleep^18^. High-throughput recording systems developed for flies include Drosophila Activity Monitor^19^, Fly Liquid-Food Interaction Counter^20^, FlyBox^21^, Whole Animal Feeding FLat^22^, DANCE^23^, and Ethoscopes^24^, many of which measure narrowly defined behaviors such as movement, feeding, aggression, or courtship. What remains less developed is a general-purpose framework that combines high-throughput recording of individual flies with comprehensive high-dimensional behavioral profile extraction.

Modern methods for analyzing behavioral recordings (Table 1) vary in two main respects. The first relates to input data modality: some methods analyze raw video frames directly, whereas others analyze body part coordinates (keypoints) obtained using markerless pose-estimation tools such as DeepLabCut^25^ (DLC) or SLEAP^26^. Converting raw video frames into keypoints can reduce dimensionality, capture salient movement features, mask irrelevant visual differences between individuals, and enable direct analysis of body part kinematics. The second distinction is the level of supervision: supervised methods require user-defined labels, whereas unsupervised methods cluster stereotyped patterns without labeled training data. Unsupervised methods, such as MotionMapper^27^, B-SOiD^28^, and Keypoint-MoSeq^29^, are powerful tools for discovering recurring structure in behavior; however mapping their native outputs onto semantic labels often requires *post hoc* interpretation, particularly in *Drosophila* where viewpoint changes (animal orientation) can fragment semantically similar actions. Conversely, supervised workflows such as JAABA^30^, DeepEthogram^31^, LabGym^32^, and SimBA^33^ provide human-readable semantic labels at the expense of manual annotation.

**Table 1:** Behavior Analysis Methods.

| Method | Input Data | Learning Style |
| --- | --- | --- |
| JAABA<sup>30</sup> | Trajectory Data and Engineered Features | Supervised |
| MotionMapper<sup>27</sup> | Raw Video/Silhouettes | Unsupervised |
| DeepEthogram<sup>31</sup> | Raw Video | Supervised |
| B-SOiD<sup>28</sup> | Keypoint-based Engineered Features | Unsupervised |
| LabGym<sup>32</sup> | Raw Video and “Pattern Images” | Supervised |
| SimBA<sup>33</sup> | Keypoint-based Engineered Features | Supervised |
| Keypoint-MoSeq<sup>29</sup> | Keypoints / Pose Dynamics | Unsupervised |
| FlyVISTA<sup>18</sup> | Keypoint-based Engineered Features | Semi-supervised |
| Autobehaver | Keypoints and Engineered Features | Supervised |

Beyond classifying individual actions, a central objective of many behavioral studies is to characterize how the overall behavioral signature of an animal is affected by experimental variables such as mutations, drugs, or environment. In many studies, identifying group-level differences is a manual and hypothesis-driven process, requiring post hoc statistical comparisons on metrics such as the frequency or duration of selected behaviors. Several studies identify group differences in more systematic ways, for example by using detailed per-animal behavior-summary vectors to discriminate between groups^30,34^, or by comparing the usage and sequencing of unsupervised behavioral syllables between groups^35,36^. Explainability methods such as SHapley Additive exPlanations (SHAP) have been proposed as a way to provide objective feature-level descriptions of behavioral differences between groups^37^, and have been applied to identify pose-derived features that contribute to behavior-label predictions^33^. However, explainable frameworks for identifying group-level differences in behavioral signatures remain less developed in *Drosophila* ethomics.

To extend interpretable, group-discriminative behavioral phenotyping to high-throughput *Drosophila* recordings, we developed Autobehaver, an end-to-end workflow that converts individual-fly videos into framewise semantic labels and high-dimensional behavioral signatures, and uses these signatures for explainable group-level phenotype discovery (Figure 1A). Autobehaver begins with movies of individual flies, from which keypoints are extracted via DLC^25^ and used to generate per-frame keypoint-derived features. These features are processed by a Transformer-based neural network to predict a Behavior and Orientation label for every frame, accounting for the perspective shifts inherent in *Drosophila* recording. These labels and keypoint-derived features are combined to generate a high-dimensional behavioral feature vector that comprehensively describes an animal’s unique behavioral signature. We then train classifier ensembles to distinguish between reference groups using these behavioral feature vectors, quantify the specific features most informative for group separation using SHAP analysis, and place intermediate phenotypes along a continuous behavioral axis. Using this framework, we recovered known circuit-driven behavioral changes, quantified age-associated decline, and resolved intermediate rescue phenotypes.

**Figure 1.**
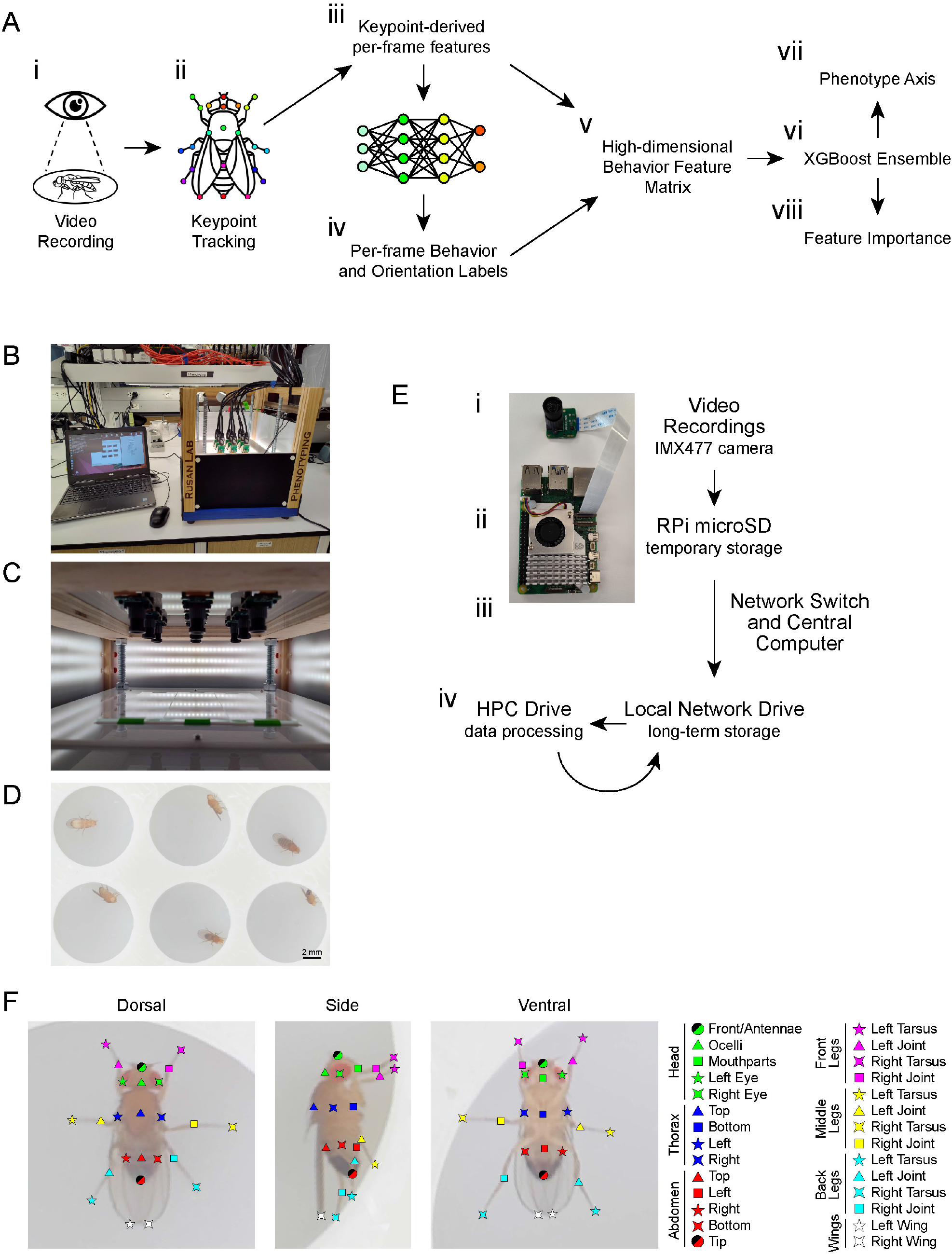
A Platform For High-throughput *Drosophila* Behavioral Recording. Overview of the Autobehaver pipeline (A): flies are recorded (i), keypoints are tracked via DLC (ii), keypoint-derived per-frame features are calculated (iii), per-frame Behavior and Orientation labels are predicted (iv), a high-dimensional behavioral feature matrix is generated (v), and an XGBoost classifier ensemble is trained (vi) to separate two groups (vii) and extract feature importance scores (viii). Photographic overview of our high-throughput recording platform highlighting the imaging box (B). Nine cameras are mounted through the top of the box, pointing downwards (C). Each camera records a bank of six 3D-printed chambers containing individual flies (D). Recordings are first captured by the cameras (E, i) and temporarily stored on the microcontroller drive (ii) before being routed through a network switch and central computer (iii) to the local network drive for long term storage and to the HPC for data processing (iv). An adult male *Drosophila* imaged from dorsal, side, and ventral perspectives shows the 30 keypoints (F) used for tracking via DeepLabCut.

## Results

### High-throughput Recording and Data Extraction for Animal Behavior

We built a recording platform (Figure 1B) with multiple low-cost cameras mounted inside of an LED-illuminated box (Figure 1C). The cameras point downwards at 3D-printed chambers containing individual flies, with six chambers per camera field of view (Figure 1D). Each camera is operated by an individual microcontroller (Figure 1E, i-ii), which are connected through a network switch to a central computer. Recordings are coordinated and data is routed to a network drive for storage and high-performance computing (HPC) cluster for processing (Figure 1E, iii-iv).

To process data for individual animals, full-frame videos were cropped into single-chamber videos and processed using a custom DLC model^25,38^ that tracks 30 keypoints on each animal (Figure 1F). Our DLC model was trained on 2,968 annotated video frames of male and female flies with a variety of distinct appearances (Supplemental Table 1), and had a median test error of 0.062 mm (Supplemental Table 2). Applying the DLC model converts a video into a matrix of x, y coordinates and likelihood values.

To prepare the keypoint matrix for neural network-based behavior prediction, we performed data corrections and feature engineering. We reasoned that prediction of behavior labels would require the network to learn how body-part positions relate to each other, and that the absolute position and angle of the fly within the chamber are irrelevant. Thus, we transformed keypoints into an egocentric coordinate system by removing absolute position and orientation, pinning the centroid at the origin and aligning the primary body axis to the positive y-axis. We then engineered features capturing whole-body movement, part-specific movement, orientation, and posture (Methods and Supplemental Tables 3-8). For each feature, we performed column-wise z-score normalization across the entire dataset. The final keypoint-derived feature matrix contained 820 feature columns (Supplemental Table 9).

### Behavior Prediction Yields High-Dimensional Per-Animal Profiles

Many widely used feature-engineered pipelines summarize temporal context within sliding windows, collapsing a span of time into aggregate statistics or spectral features and thereby discarding information about the precise timing of micro-movements. Transformer architectures^39^, by contrast, model ordered sequences directly, making them attractive for learning subtle temporal structure from behavioral data. We developed a custom Transformer-based architecture to predict behavior labels (Figure 2A). The model input was a sequence of rows/frames from the keypoint-derived feature matrix, consisting of a focus frame to be labeled which is flanked by past and future frames for temporal context. This sequence of frames was processed via a Transformer encoder to generate a latent representation which was passed through two prediction heads to predict Behavior and Orientation labels. The model was trained using a custom dataset comprising ~2.7 hours of manually labelled frames (Supplemental Table 10) derived from video clips of over 400 flies (Supplemental Table 11). Each frame was labelled with one of ten possible Behavior labels (Standing, Walking, Scrambling, Jumping, Head Grooming, Front Leg Grooming, Side Leg Grooming, Wing Grooming, Abdomen Grooming, and Back Legs Grooming; Supplemental Figure 1A and Supplemental Video 1) and also one of five possible Orientation labels (Floor, Ceiling, Wall-Side, Wall-Up, and Wall-Down; Supplemental Figure 1B).

**Figure 2.**
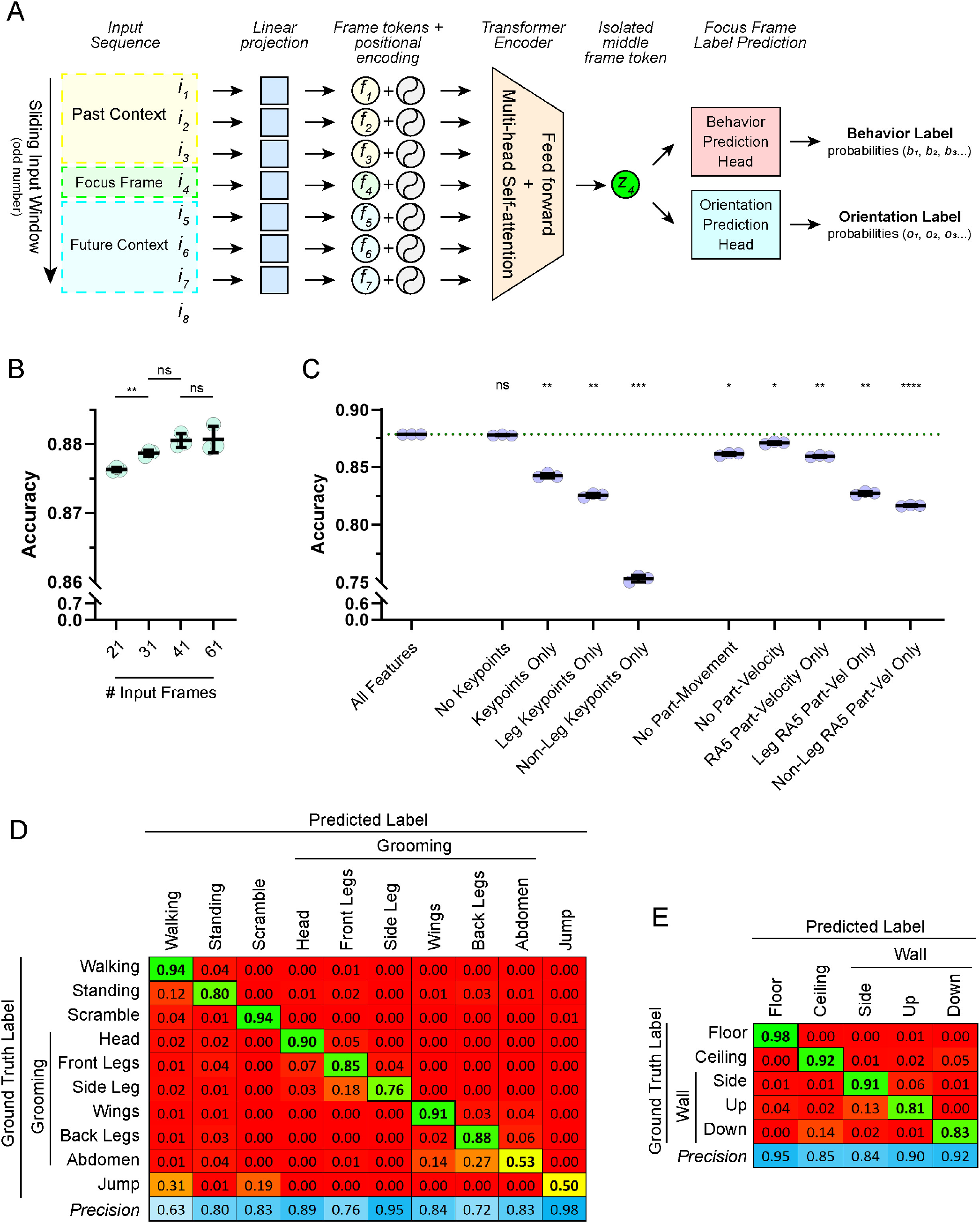
Autobehaver Predicts *Drosophila* Behavior and Orientation. (A) Autobehaver model architecture. An odd number of frames (*i*; 7 shown for simplicity) forms the input sequence, consisting of a focus frame (*i*_*4*_) to be labelled, along with an equal number of past (*i*_*1*_-*i*_*3*_) and future (*i*_*5*_-*i*_*7*_) frames for temporal context. Each input frame passes through a linear input layer to become a frame token (*f*), a learned vector representation of that frame. Positional encodings are added to the frame tokens, and the sequence is passed through a Transformer encoder. The transformed middle frame token (*z*_*4*_) is isolated as the latent representation and passed through two prediction heads to predict Behavior and Orientation labels for the focus frame. To generate predictions for a full movie, the input window slides forward one frame at a time so that each frame is treated as the focus frame for one forward pass. Model accuracy improved as the number of input frames increased, up to 41 frames (B); see Supplemental Figure 2 for full details on architecture tuning. Dataset ablation experiments (C) show the average model accuracy obtained with dataset configurations, highlighting the differences in the information content of leg vs non-leg keypoints and part-specific velocity features; see Supplemental Figure 3 for full details on dataset ablations. Confusion matrices for the “All Features” models shows the recall (diagonal, bold outlines) and precision (bottom row) for Behavior (D) and Orientation (E) predictions. Color scale for recall ranges from red (0) to green (1), and the color scale for precision ranges from white (0.5) to medium blue (1). Statistical comparisons were made using Welch’s ANOVA with Dunnett’s T3 post-hoc test for all-to-all pairwise comparisons (B) or comparisons against “All Features” (C). **** p < 0.0001; *** p < 0.001; ** p < 0.01; * p < 0.05; ns, not significant.

We evaluated training hyperparameters and architectural configurations to maximize macro F1 and accuracy on our validation dataset. Performance peaked with a learning rate of 1 × 10^−4^ (Supplemental Figure 2A) and an embedding dimension of 1024 (Supplemental Figure 2B). The best-performing models were trained using 41 input frames (Figure 2B, Supplemental Figure 2C), a single encoder layer (Supplemental Figure 2D), 16 to 64 attention heads (Supplemental Figure 2E), and using a combination of middle-frame representation, sinusoidal positional encodings, and a multi-layer perceptron prediction head (Supplemental Figure 2F).

The full keypoint-derived feature matrix contained 820 features. To determine whether the model relies on information from each feature category, we performed dataset ablations by training with a common model architecture while either dropping one or more feature subsets (Supplemental Figure 3A) or retaining only a subset of features (Supplemental Figure 3B). For example, removing all body-part-specific movement features led to 1.7% drop in validation accuracy compared to training on the full dataset (Figure 2C). Removing most feature categories caused only minor fluctuations in performance (Supplemental Figure 3A), with the exception of part-specific movement features (with part-specific velocity being the most informative subset). Retaining only the keypoints (either with or without likelihoods) provided surprisingly good performance (3.6% and 4.3% drop in validation accuracy, respectively), with the legs contributing more than other keypoints. Similarly, retaining only part-specific movement, part-specific velocity, or part-specific velocity calculated from a five frame rolling average window, yielded very good performance (1.9% drop in validation accuracy in each case), with the legs again contributing more than other keypoints. Confusion matrices for the validation set show that the best-performing model architecture trained on the full keypoint-derived feature matrix had difficulty with Abdomen Grooming (which was often confused with Back Leg Grooming and Wing Grooming) and Jump (often confused with Scrambling or Walking; Figures 2D and 2E). These lower-accuracy predictions were likely due to similar limb dynamics of grooming behaviors, and the very brief and rare nature of jumping, respectively. Additional training data would likely improve label accuracy. We trained an ensemble of three final models (Supplemental Table 12) using the entire training set without a validation split, saving the checkpoint from the final epoch. During inference, we performed three forward passes per model with test-time augmentation (minor noise) to mitigate model-specific errors and obtain a more robust final prediction. Performing inference predicts a Behavior (B) label and an Orientation (O) label for each frame, and the combined frame-wise state is herein referred to as the BxO class (see Supplemental Video 2 for example inference).

To compare behavior across individual animals and groups, we used the predicted BxO labels and keypoint-derived feature matrix to derive high-dimensional feature matrices that comprehensively describe an animal’s unique behavioral signature. Consecutive frames with the same BxO label were grouped into bouts, and these bouts were used to organize multiple categories of summary features. These included time-budget and bout-summary features, behavioral-transition features, body measurements, postural, spatial, and sleep features, as well as an extensive set of kinematic features computed by first calculating per-bout summary statistics and then calculating mean and standard deviation across all bouts of a given BxO class. These features were aggregated per-animal to generate a high-dimensional behavioral feature vector containing 9,566 total features that captured the time spent and the characteristic posture and movement associated with each BxO class, and formed the basis for subsequent group-level analyses. See Methods and Supplemental Tables 13 and 14 for full details.

### Segmentation and Analysis of Discrete Behavioral and Morphological Features

To validate whether these behavioral feature matrices were biologically-grounded, we analyzed flies with easily identifiable behavioral characteristics. We used the bipartite GAL4-UAS system^40^ to express the temperature-sensitive Transient receptor potential Ankyrin 1 (*UAS-dTrpA1*) cation channel^41^ in two subsets of neurons: one subset driven by *29E04-GAL4*^42^ and another driven by *GH146-*GAL4. Flies were raised at 18°C to prevent dTrpA1 activation, and then transferred to the recording chamber, which was maintained at 31 ± 1°C (Supplemental Figure 4), to trigger neuronal depolarization via dTrpA1. Flies carrying *UAS-dTrpA1* but no GAL4 driver were used as controls. We then recorded these animals and analyzed their behavior.

The *29E04-GAL4* line contains an enhancer fragment from the intron of *Netrin-B* and was characterized as a potent driver of wing-grooming behavior when paired with dTrpA1^43^. When analyzed by Autobehaver, flies expressing *29E04-GAL4*-driven *dTrpA1* exhibited a 3.8-fold increase in time spent Wing Grooming (Figure 3A, pink shading). Autobehaver also captured this increase via other Wing Grooming-related features: for example, increased bout frequency (Figure 3B), longer average bout duration (Figure 3C), and shorter average time between bouts (Figure 3D). Next, we tested Autobehaver’s ability to quantify broad shifts in arousal and locomotor states using *GH146-GAL4*, which is expressed in the ~90 olfactory projection neurons (OPNs) and the anterior paired lateral neuron of each hemisphere^44,45^. Flies with *GH146-GAL4*-driven *dTrpA1* showed a striking increase in the proportion of time spent Walking and a corresponding decrease in Standing, consistent with increased arousal from acute OPN activation (Figure 3A). Autobehaver also identified a significant reduction in sleep bouts (defined as Standing for ≥ 3 minutes; Figure 3E), a decrease in the time between Walking bouts (Figure 3F), and a decrease in average Walking speed (Figure 3G). Together, these experiments show that Autobehaver can quantify both the induction of specific behaviors and changes associated with global shifts in arousal state.

**Figure 3.**
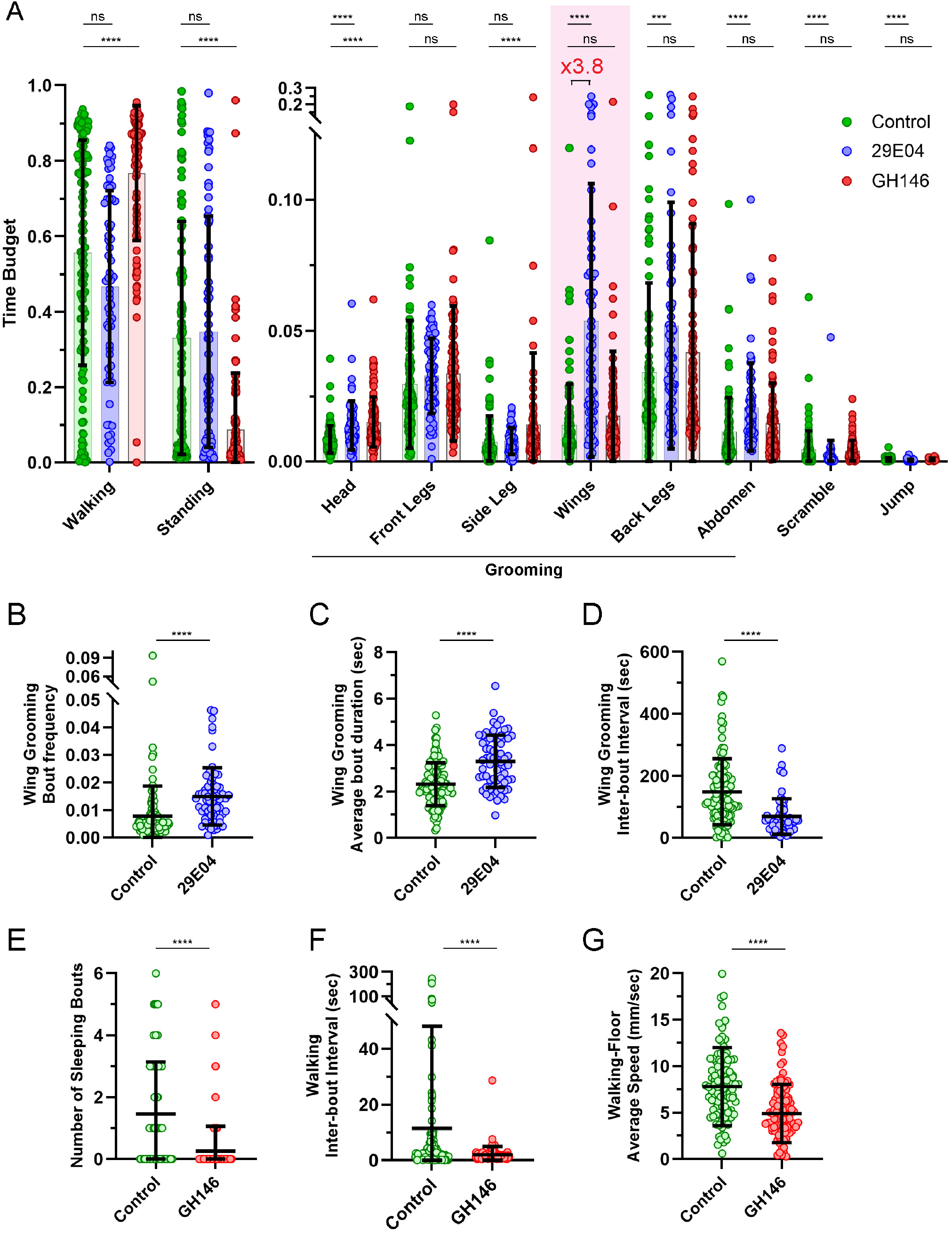
Autobehaver Detects Discrete Behavioral Differences. Per-animal behavioral time budgets (A) of Control (*UAS-TrpA1* with no driver; green), *29E04-GAL4* > dTrpA1 (blue), and *GH146-GAL4* > dTrpA1 (red), with statistical significance assessed via Mann-Whitney test with Holm-Bonferroni multiple comparison correction. Per-animal wing-grooming bout frequency (B), average bout duration (C), and inter-bout interval (D) all show significant differences between Control and *29E04-GAL4* > dTrpA1. Compared to Controls, *GH146-GAL4* > dTrpA1 show significantly lower number of sleeping bouts (E), decreased inter-bout interval (F), and lower average Walking speed while on the Floor (G).

Aging in *Drosophila* is associated with progressive locomotor decline^46^. We tested whether Autobehaver could detect and quantify the physiological differences among flies aged 4, 30, or 60 days post-eclosion. First, we found a progressive decrease in the proportion of time spent Walking, and a corresponding increase in the proportion of time spent Standing as the flies aged (Figure 4A). Additionally, we found that flies spent increasingly more time on the Floor and less time on the Ceiling with age (Figure 4B), suggesting a decline in climbing ability^47^. Investigation of kinematic features revealed that 60-day-old flies had significantly slower Walking velocity compared to younger cohorts (Figure 4C), and that flies showed a reduction in maximum Walking acceleration with age (Figure 4D). Altogether, these results show that Autobehaver can quantify gradual multidimensional shifts in behavioral vigor such as those seen in aging.

**Figure 4.**
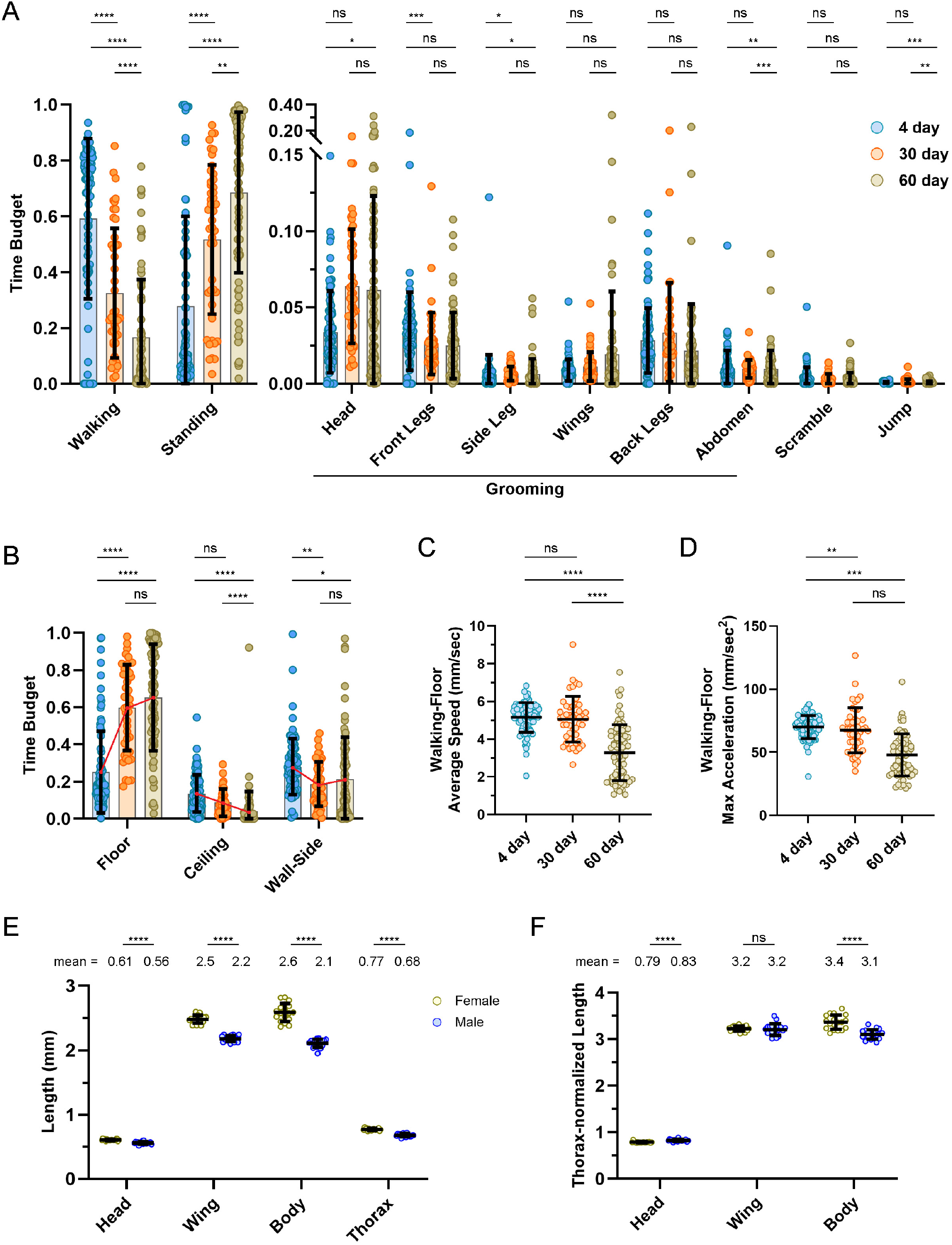
Autobehaver Reveals Age- and Sex-related Differences. Per-animal behavioral (A) and orientation (B) time budgets of male control flies aged to 4 days (blue), 30 days (orange) and 60 days (tan), with statistical significance assessed via Mann-Whitney test with Holm-Bonferroni multiple comparison correction (total comparisons combined for A and B). Compared to 4-day-old and 30-day-old flies, 60-day-old flies had significantly reduced speed while Walking on the Floor (C). Compared to 4-day-old flies, 30-day-old and 60-day-old flies had significantly reduced maximum acceleration while Walking on the Floor (D). Comparison of female (yellow) and male (blue) body size features showed a significant reduction in male size across all measurements (E). When body size features were normalized by thorax width, male heads were proportionally slightly larger, wing length was not different, and female body length was proportionally longer (F). Statistical comparisons were made using Mann-Whitney test with Holm-Bonferroni correction for multiple comparisons. **** p < 0.0001; *** p < 0.001; ** p < 0.01; * p < 0.05; ns, not significant. C.P., mean Class Prediction; P.S., mean Phenotype Score.

### Automatic Extraction of Body-size Measurements

Leveraging keypoint accuracy, we captured morphological measurements from frames where the animal maintained a standard pose, such as Standing on the Floor. As expected, male flies were significantly smaller than female flies across all size measurements, including head width, wing length, body length, and thorax width (Figure 4E). Insect body size scales proportionally, and thorax width is often used as a size normalizer^48^. We therefore normalized the measurements from each individual fly by its thorax width and compared relative body proportions. After normalizing measurements by thorax width, we found that male head width was slightly larger, wing length was equivalent, and female body length was slightly longer (Figure 4F). Thus, Autobehaver can complement behavior classification with automated morphometric analysis, providing an additional tool for identifying phenotypes involving structural changes.

### Classifier-based Predictions as a Proxy for Behavioral Similarity

We next asked whether per-animal behavioral feature vectors could be leveraged to reveal group-defining behavioral signatures. Although behavior varies across individual flies, members of the same group (e.g., genotype or treatment) should share behavioral signatures that distinguish them from other groups. Based on this premise, we trained XGBoost classifiers, a gradient-boosted ensemble of decision trees^49^, to predict the group identity of individual flies from a matrix of behavioral feature vectors (Figure 5A). Our downstream goal was to use these classifiers both for prediction and to identify features most informative for separating groups.

**Figure 5.**
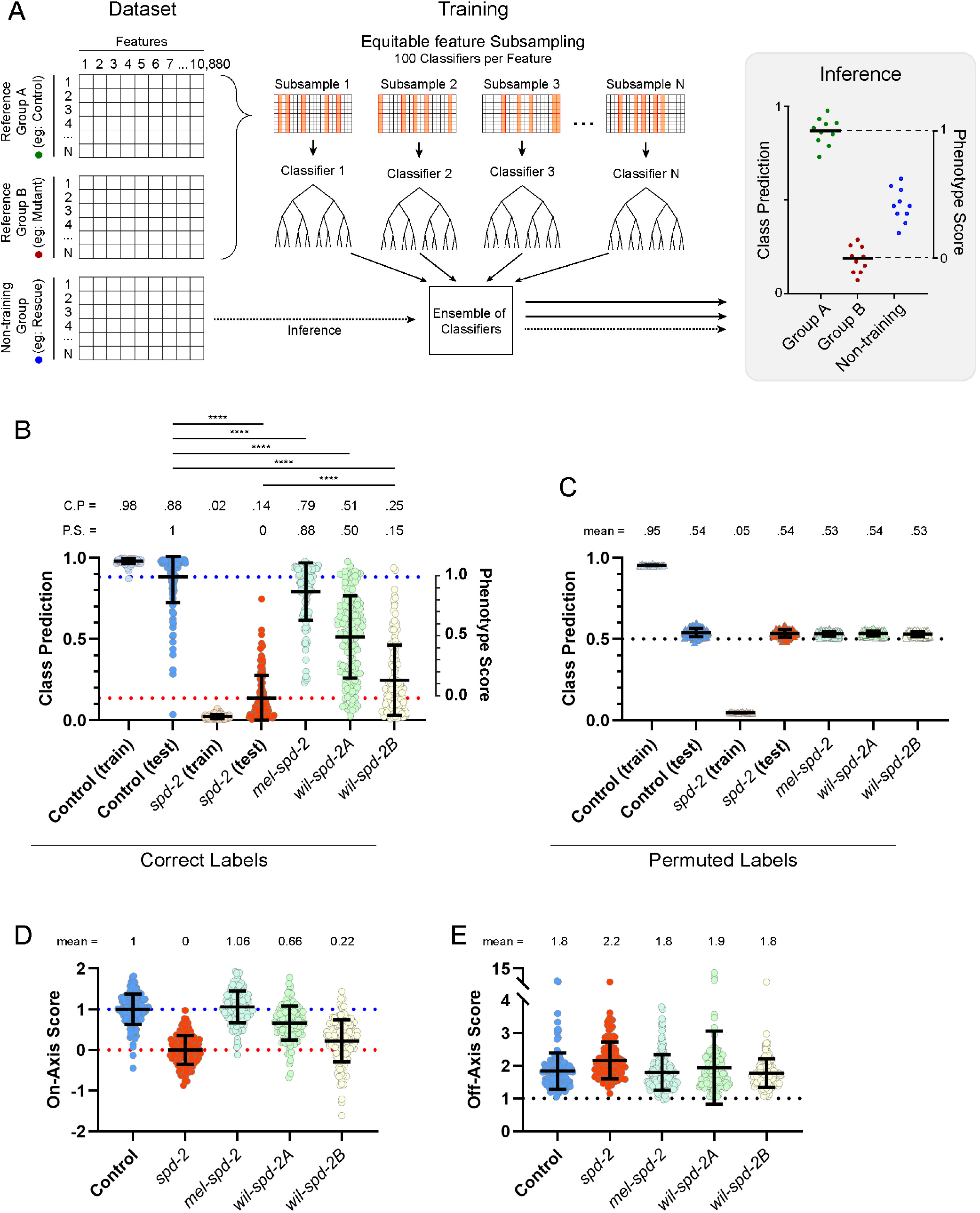
Behavior-based Classifiers Reveal Intermediate Behavioral States. Overview of behavioral classifier training and prediction (A). The full dataset comprises individual flies (rows) from multiple groups, each with 9,566 features (columns). A pair of reference groups (eg: Control and Mutant) is used to train an ensemble of XGBoost classifiers. During training, each model is fit on a random subset of the total features, resulting in an ensemble containing thousands of classifiers that together produce an aggregate class prediction. We linearly rescale the class predictions between the mean predictions of the reference groups to produce a Phenotype Score for interpreting behavioral similarity. Training a classifier ensemble to distinguish between 15-day-old control and *spd-2* mutants (B) produced strong discriminative power (blue and orange dotted lines) and a class prediction separation of 0.74, enabling quantification of intermediate behavior phenotypes. Applying the classifier to *spd-2* mutants carrying different transgenes showed that *mel-spd-2, wil-spd-2A* and *wil-spd-2B* have varying levels of behavioral rescue and intermediate PS. Training a classifier ensemble on individuals with permuted labels (C) results in no discriminative power (average class prediction for non-training individuals = 0.53). The full behavioral feature matrix was projected into principal component space to define an axis separating the centroids of control and *spd-2* individuals, resulting in on-axis (D) and off-axis (E) components for each individual, reported in the same unitless scale. On-axis values indicate each fly’s position along the line connecting the average positions of the two reference groups in PCA space; off-axis values indicate each fly’s distance from that line. Statistical comparisons were made using Mann-Whitney test with Holm-Bonferroni correction for multiple comparisons. **** p < 0.0001. C.P., mean Class Prediction; P.S., mean Phenotype Score.

An XGBoost model trained on the full behavioral feature matrix might concentrate predictive weight on a small number of highly informative features, limiting its usefulness for feature-importance discovery. To distribute model attention more broadly, we trained a multi-round classifier ensemble using five-fold cross-validation, where in each round the feature space was partitioned into random subsets of 100 features and one model was trained per fold for each subset. We designed the ensemble so that each feature appeared in ~100 different random subsamples. This repeated random feature subsampling was done so that downstream feature-importance analysis could capture signal distributed more broadly across the behavioral feature space.

For each feature subset, out-of-fold performance across the five folds was pooled to generate a subset-level performance weight, such that subsets outperforming a permuted-label null contributed more strongly to the final prediction. We then aggregated the raw margins from all classifiers in logit space using these subset-level weights and converted the result to a class prediction between 0 and 1. To calibrate these predictions, we linearly rescaled them using the mean out-of-fold ensemble predictions of the two reference groups, producing a Phenotype Score (PS; 0 to 1) that reflects an individual’s position along the classifier-defined discriminative axis.

To validate whether the PS could accurately capture the gradations between biological states, we applied this framework to *spd-2*, a gene important for the microtubule nucleation function of centrosomes and proper mitotic spindle formation. A previous study reported that *spd-2* mutants exhibit uncoordinated movement^50^. Our previous work investigated the functions of *spd-2* from *Drosophila melanogaster* and the two paralogous genes from *Drosophila willistoni, spd-2A* and *spd-2B*^51^. In that study, we showed that a *mel-spd-2* transgene strongly rescued mitotic centrosomal activity in the larval brain, a *wil-spd-2A* transgene moderately rescued, and a *wil-spd-2B* transgene did not rescue. We hypothesized that if behavioral output reflects underlying cell biology, then behavioral signatures from individuals expressing these rescue transgenes should yield Phenotype Scores that mirror the degree of cell biological rescue.

To test this hypothesis, we collected control and *spd-2* flies, recorded behavior, applied keypoints via DLC, generated keypoint-derived feature matrices, predicted BxO labels, calculated behavioral feature vectors for individuals, used the resulting matrix of behavioral feature vectors to train a classifier ensemble to separate control and *spd-2*, and obtained class predictions which were rescaled into a PS. We then used the classifier ensemble to process behavioral feature vectors from the three transgenes (Figure 5B). We found that *mel-spd-2* caused a strong shift toward control (PS: 0.88), *wil-spd-2A* caused a ~50% shift toward control (PS: 0.50), and *wil-spd-2B* remained near the *spd-2* null flies (PS: 0.15), thus mirroring the cell biology^51^. These results demonstrate that the classifier ensemble can place intermediate phenotypes along a continuous behavioral axis defined by two reference groups.

To test whether intermediate class predictions reflected genuine genotype-linked structure in behavioral feature space, rather than an artifact of the model, we performed a label-permutation experiment by randomly shuffling the reference group labels and retraining the classifier ensemble using an identical pipeline. With the genotype-behavior relationship broken, the permuted classifier ensemble was unable to produce decisive predictions for out-of-fold validation individuals and non-reference groups, producing class predictions near 0.53 (Figure 5C). As an independent, unsupervised check, we also projected the behavioral feature matrix into PCA space, which can be viewed as a simplified map of overall behavioral similarity. We then defined a control-mutant axis as the line connecting the average control and *spd-2* positions. Each fly’s on-axis value is its position along this line, whereas its off-axis value is its distance away from the line, both reported in the same unitless scale. The rescue genotypes showed a similar ordering along the on-axis dimension (Figure 5D), indicating that they occupy real intermediate behavioral states rather than positions created by the classifier. Off-axis distances remained substantial (Figure 5E), indicating additional behavioral variation outside this main control-mutant continuum. These results indicate that the graded Phenotype Scores arise from reproducible, group-associated behavioral signatures that generalize beyond the reference individuals.

### Revealing Group-specific Behavioral Features Via Importance Analysis

We next developed a feature-importance analysis to identify the behavioral features most informative for distinguishing between two reference groups (genotype, treatment, etc.). In XGBoost, predictions are built through sequential decision nodes, each of which selects a feature and threshold that improves classification (Figure 6A, panel i). We quantified feature importance within our multi-round subsampling classifier ensemble using SHapley Additive exPlanations^52^ (SHAP), which assigns each feature a signed contribution to an individual prediction (Figure 6A, panel ii). Before aggregation across models, SHAP values from each feature subsample were weighted by subset-level out-of-fold predictive performance so that reliable subsamples contributed more strongly to the final importance estimates. We summarized SHAP-based importance in two ways: group-specific mean SHAP, defined as the mean signed out-of-fold SHAP value within each reference group; and global importance, defined as the mean absolute weighted out-of-fold SHAP value across individuals from both reference groups, after subtraction of the permutation-derived SHAP baseline (Figure 6A, panel iii). These summaries identify the features that separate the reference groups, and encode the magnitude and direction of their contributions to classification.

**Figure 6.**
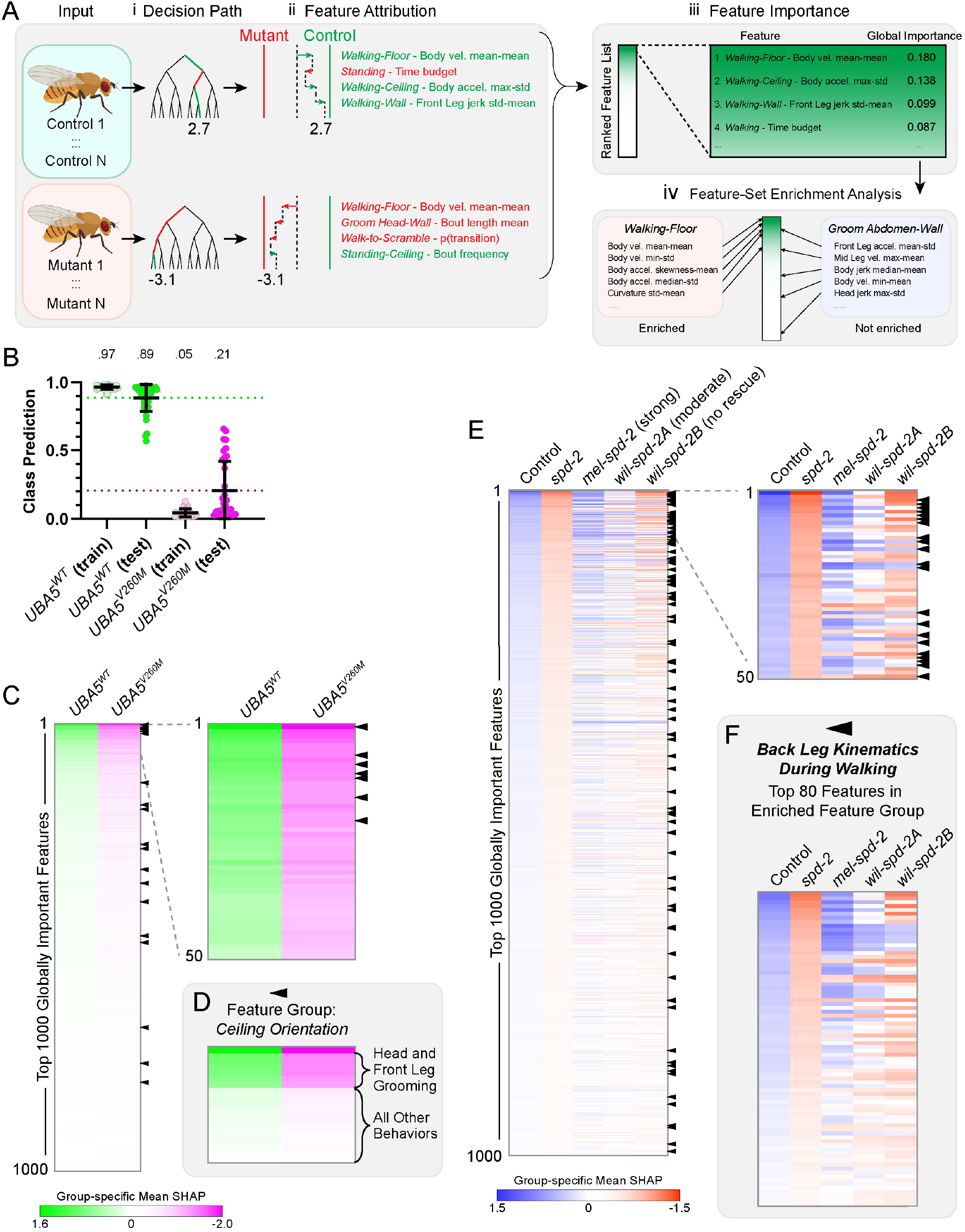
Autobehaver Reveals Group-specific Behavioral Differences. Diagram showing feature-importance analysis (A). Data from individual flies from two reference groups are used to train an XGBoost classifier ensemble, which is then used to generate a prediction for each individual (i). We calculate a SHAP value for the feature for each individual (ii), and then aggregate them into a single group-specific mean SHAP value for each of up to 9,566 behavioral features (iii), where the magnitude indicates the importance and the sign indicates which direction that feature tends to influence the prediction. Groups of features can be analyzed via feature set enrichment analysis (FSEA) to identify feature sets that are highly enriched near the top of the ranked list and thus contribute strongly to the phenotypic profile of that group (iv). A classifier ensemble discriminates between *UBA5*^*WT*^ and *UBA5*^*V260M*^ (B). Barcode plots show group-specific mean SHAP values for *UBA5*^*WT*^ (green, positive direction) and *UBA5*^*V260M*^ (magenta, negative direction) for the top 1000 and top 50 globally important features (C). FSEA identifies 12 enriched feature groups for *UBA5*, including the feature group containing bout summary statistics for Ceiling-oriented bouts; black pointers highlight specific features from this group along the main barcodes in 6C, and the top 20 Ceiling-oriented bout features are shown as a separate barcode (D). Barcode plots show group-specific mean SHAP values for control (blue, positive direction) and *spd-2* mutants (vermillion, negative direction) for the top 1000 and top 50 globally important features (E). Group-specific mean SHAP values for *mel-spd-2, wil-spd-2A* and *wil-spd-2B*, corresponding to strong, moderate, and no rescue, respectively, are in the rightmost three columns, indicating the direction each feature influences the prediction. FSEA identifies enriched feature groups for *spd-2*, including the feature group describing back leg kinematics during Walking bouts; black pointers mark the features from this group along the main barcodes in 6E, and the top 80 features from this group are shown as a separate barcode (F).

To identify patterns among the most important features, we first inspected the top 50 features ranked by global importance. Because thousands of individual features are difficult to interpret by eye, we applied Feature Set Enrichment Analysis (FSEA), adapted from gene set enrichment analysis^53^ and implemented using the GSEApy Python package^54^, on the SHAP-ranked feature list to systematically assess whether predefined feature groups were overrepresented among the highest-ranked features (Figure 6A, panel iv). FSEA scans down the ranked list and assigns higher enrichment scores to groups whose members appear disproportionately early. Feature groups were generated programmatically by grouping related metrics, such as body part-specific kinematics, time budgets, transition probabilities, or sleep measures, within specific BxO contexts. This collapsed many individual features into more interpretable behavioral themes; for example, one group contained all front leg kinematic features measured during Head Grooming-on-Floor bouts (Supplemental Table 14).

To assess whether feature-importance analysis could recover biologically meaningful patterns, we examined a humanized fly model in which human *Ubiquitin-like Modifier Activating Enzyme 5* (*UBA5*) replaces the fly gene^55^. *UBA5* is the E1 activating enzyme for UFMylation, a post-translational modification involved in protein homeostasis and endoplasmic reticulum stress regulation^56,57^. In humans, hypomorphic *UBA5* mutations are linked to multiple conditions, including Developmental and Epileptic Encephalopathy 44 (DEE44), which is characterized by encephalopathy, developmental delay, abnormal movement, and seizures^58,59^. In *Drosophila, uba5* knockdown causes wing postural and climbing defects^60^, whereas a *uba5* null mutation is embryonic lethal^55^. Expressing a wild-type (WT) human *UBA5* transgene in the *uba5* mutant background rescues lethality with no detectable defects, whereas expressing DEE44-associated alleles in the *uba5* mutant background results in adults with climbing and seizure defects^55^.

To test whether Autobehaver could detect these expected behavioral changes, we analyzed recordings of 30-day-old male *uba5* null flies expressing either *UBA5*^*WT*^ or the severe DEE44-associated allele *UBA5*^*V260M*^ and trained a classifier ensemble to discriminate between them (Figure 6B). A permuted-label classifier failed to recover meaningful separation (Supplemental Figure 5A), and an unsupervised PCA of the full behavioral feature matrix still separated *UBA5*^*WT*^ and *UBA5*^*V260M*^ (Supplemental Figure 5B,C), indicating that the classifier was using genuine group-associated behavioral structure rather than overfitting. We calculated group-specific mean SHAP and global importance scores for all features and ranked features by global importance (Figure 6C).

To identify the most informative features used to distinguish *UBA5*^*V260M*^ from *UBA5*^*WT*^, we began by examining the top 50 globally important features (Figure 6C; Supplemental Table 16). Ceiling-related features dominated this set (22/50 features). Inspection of the raw feature data revealed that *UBA5*^*V260M*^ individuals had shorter Ceiling bout duration, narrower leg posture, reduced walking kinematics, less time spent Head Grooming and Front Leg Grooming, and lower probability of transitioning from Walking to Standing while on the Ceiling (Supplemental Figure 5D-K), consistent with previously reported climbing defects^55^. Scrambling-related features were also prominent (8/50; bouts in which flies fall onto their backs and attempt to right themselves), reminiscent of the reported seizure-sensitivity phenotype^55^. FSEA of the ranked list identified 12 significantly enriched feature groups (Supplemental Table 17), including Orientation bout summary features that captured reduced Ceiling-oriented behaviors, Scramble-related bout statistics and kinematic features, feature groups of bout statistics for Head Grooming, Front Leg Grooming, and Side Leg Grooming categories, and the feature group of body measurements (Supplemental Table 17). Within the Ceiling-Orientation feature group, the strongest group-specific mean SHAP signals were concentrated in Ceiling bout duration and in the proportion of Ceiling time spent Head Grooming and Front Leg Grooming, whereas other Ceiling-associated features contributed more weakly (Figure 6D). This pattern suggests that *UBA5*^*V260M*^ flies may be particularly impaired in upside-down behaviors that require lifting the front legs from the substrate. We also examined the raw feature values for the body measurement feature group and found that the *UBA5*^*V260M*^ flies had significantly reduced head, thorax, and abdomen width, but no difference in body or wing length compared to *UBA5*^*WT*^ controls (Supplemental Figure 5L-P). Together, these analyses implicated climbing-related locomotor defects, Scrambling and Grooming phenotypes, and subtle size differences, highlighting biologically meaningful group differences recovered by Autobehaver.

To better understand the basis of intermediate predictions, we revisited the *spd-2* transgenic rescue series and performed SHAP-based analysis (Figure 6E; Supplemental Table 18). Inspection of the top 50 globally important features suggested that back leg kinematics during Walking bouts were especially prominent (21/50 features), with additional representation of Scrambling-related features (6/50 features; Supplemental Table 18). FSEA revealed 5 enriched feature groups, including bout-statistics groups for Walking, Standing, and Side Leg Grooming, the feature group containing bout statistics and kinematics of Scrambling, and the feature group containing back leg kinematic features during Walking (Supplemental Table 19). These globally enriched feature groups tracked the rescue continuum: for example, we examined group-specific mean SHAP values for back leg kinematic features during Walking across the three transgenic lines and found that they followed the strong, moderate, and no rescue series (Figure 6F). Inspection of the underlying feature values suggested that *spd-2* mutants exhibit more variable back leg movement dynamics during Walking, with higher within-bout variability and higher peak velocity, acceleration, and jerk compared to controls (Supplemental Figure 6A-H). Together, these analyses show that Autobehaver can identify both specific behavioral features and feature groups most informative for distinguishing reference groups, while also helping interpret where intermediate phenotypes fall along the resulting behavioral axis.

### Comparison of Subsampling and Full-feature XGBoost Feature-Importance Strategies

Our feature-importance framework uses a complex subsampling approach to train an ensemble of XGBoost classifiers, in which each model is trained on only a small random subset of the full feature space. We designed this subsampling strategy to reduce competition among features, so that informative but weaker features would contribute to the final importance profile rather than being masked by the most strongly predictive features. By contrast, a standard full-feature XGBoost model is expected to concentrate importance on a smaller set of dominant features. We therefore compared our subsampling approach with an ensemble of full-feature XGBoost models to test whether subsampling broadened feature-importance estimates as expected. Classification performance was broadly similar between the two approaches, with only small dataset-dependent differences in area under the curve and F1 score (Supplemental Figure 7A,B).

This indicates that discriminative information is distributed across many features, because classifiers trained on smaller feature subsets still effectively separated reference groups. To assess how concentrated or broadly distributed the importance signal was, we plotted the cumulative fraction of total global importance captured as features were added from highest to lowest rank. Full-feature XGBoost rose sharply and plateaued early, indicating that importance was concentrated in a relatively small number of dominant features. By contrast, the subsampling approach accumulated importance more gradually across the ranked list, indicating that importance was distributed across a broader range of features (Supplemental Figure 7C). This result is consistent with the design goal of subsampling, namely to reduce feature competition and allow informative but weaker features to contribute to the final importance profile.

We next asked whether the feature rankings were reproducible across three independent runs of each method. We assessed this in two complementary ways: Jaccard overlap measured whether repeated runs recovered the same top-k features, whereas Spearman rank correlation measured whether those features appeared in a similar order. Full-feature XGBoost was highly consistent for the top-ranked features, but this consistency declined as larger portions of the ranked list were considered. In contrast, the subsampling approach maintained high overlap and rank-order agreement across a broader span of the ranked list (Supplemental Figure 7D,E). Between methods, overlap and rank-order agreement were moderate and dataset-dependent, indicating that the two approaches produced overlapping but non-identical ranked feature lists (Supplemental Figure 7D,E).

Subsampling also improved downstream Feature Set Enrichment Analysis. In both datasets, the subsampling approach consistently recovered a core set of positively enriched feature groups across repeated runs (Supplemental Tables 17 and 19), whereas full-feature XGBoost did not recover any significantly enriched groups. This likely reflects the broader and more reproducible distribution of feature importance produced by subsampling, because enrichment analysis depends on related features appearing together near the top of the ranked list. Together, these results indicate that the main benefit of subsampling was a broader and more reproducible feature-importance profile that better supported biologically interpretable enrichment analysis.

## Discussion

Here we present Autobehaver, our automated high-throughput behavioral phenotyping pipeline. We integrated modular, low-cost hardware with Transformer-based sequence modeling of DLC-derived keypoints to generate bout structure and extract 9,566 behavioral features per animal. We then trained XGBoost classifier ensembles to distinguish between two reference groups, enabling (i) continuous quantitative scoring of intermediate behavioral profiles, and (ii) interpretable feature attribution using SHAP analysis to prioritize individual features defining group-specific behavioral signatures.

We validated Autobehaver’s utility by analyzing several distinct datasets spanning discrete, gradual, and intermediate phenotypes. First, we used Autobehaver to quantify circuit-specific shifts in time budgets and bout statistics using temperature-sensitive dTrpA1 to activate specific neural circuits (Figure 3A-G). Second, we used Autobehaver to capture progressive decline in locomotor vigor and climbing in an aging series (Figure 4A-D). We also used Autobehaver to extract keypoint-based morphometrics as an orthogonal phenotypic readout (Figure 4E,F). Finally, we used Autobehaver to quantify intermediate behavioral phenotypes falling between two reference groups by characterizing a series of transgenic lines with different levels of phenotypic rescue (Figure 5B,C).

To summarize intermediate behavioral signatures, we defined a Phenotype Score (PS) relative to the two reference groups. Importantly, PS reflects similarity along the discriminative behavioral axis learned by the classifier ensemble. It is therefore not informative about behavioral features that contribute little or nothing to reference-group separation, nor is it an absolute or cross-experiment comparable measure of phenotypic severity. To interpret the features underlying this axis, we combined SHAP-based feature-importance analysis and FSEA to prioritize individual features and feature groups associated with group separation and intermediate phenotypes (Figure 6D, 6E). Thus, Autobehaver provides a framework for quantifying behavioral similarity while highlighting candidate behavioral modules for follow-up study.

Our first-generation recording platform uses modular, low-cost camera-microcontroller units with high spatial and temporal resolution and moderately high throughput. The current system ran at ~32°C, which was used for acute neuronal activation via the heat-sensitive dTrpA1 channel, but this temperature is above the standard physiological temperature for *Drosophila*. Future design iterations will implement temperature control. Notably, the elevated temperature did not prevent the identification of group-defining behavioral profiles; rather, elevated temperature or other environmental stressors may amplify group-specific differences in behavioral signatures. A second limitation of the current design was the small chamber size (9 mm diameter, 2.5 mm height), which restricted flight but was required to maintain depth of field and enable parallel high-throughput recording. In the current design, all groups were collected, aged, handled, and recorded in parallel, and genotypes/sexes were staggered across chambers and rotated across trials so that each condition was sampled across camera and chamber positions. Overall, the current system provided a simple and low-cost, yet robust and reproducible recording platform.

Dynamic behaviors are defined by coordinated patterns of motion over time. Thus, we used a Transformer encoder architecture to integrate a one-second temporal window and predict Behavior and Orientation labels. Combining the ability of a Transformer to capture temporal structure with a human-designed feature space ensured that Behavior and Orientation predictions were grounded in interpretable, biologically meaningful patterns. For *Drosophila*, explicit Orientation labels are useful because floor, ceiling, and wall contexts alter both the visual appearance of behavior and its biological interpretation. Future work could hybridize the supervised label ontology with unsupervised pose-dynamic discovery methods such as keypoint-MoSeq^29^, which can recover modules aligned with human annotations while also revealing structure beyond predefined labels. The Autobehaver codebase is modular and can be adapted to alternative keypoints or engineered features.

Several limitations of the current Autobehaver implementation warrant consideration. First, Autobehaver is inherently constrained by its user-defined vocabulary of Behaviors and Orientations. Rare or ultra-brief behaviors such as Jumping, which typically spans 1–3 frames, are challenging to learn given limited training examples (Figure 2F). Expanding the label repertoire is straightforward, but requires additional manual annotation and retraining of the Transformer network. SHAP- and FSEA-based feature prioritization should be viewed as hypothesis-generating: these analyses identify features and feature groups associated with group discrimination, but do not by themselves establish causal mechanisms. Our current FSEA feature groups are predefined and generated programmatically from broad feature categories, which may not capture the true biological structure of coordinated behavioral change. Future work could define feature groups using empirical co-variation or context-specific priors to produce enrichment analyses that are more biologically grounded. Because the recording setup was intentionally standardized (constant chamber geometry, optics, and lighting), substantially changing the recording parameters would likely require retraining or fine-tuning. While we focus here on *Drosophila*, Autobehaver is adaptable to other organisms and recording setups.

Existing frameworks have established the value of automated behavioral phenotyping in *Drosophila* and other systems. These include early *Drosophila* ethomics pipelines that represented behavior as statistical vectors and recovered sex- or genotype-linked structure from video-derived data^34^, supervised classifiers such as JAABA^30^, DeepEthogram^31^, LabGym^32^, and SimBA^33^, related temporal-context classifiers^61^, and more recent *Drosophila* deep-phenotyping platforms such as FlyVISTA^18^. Unsupervised approaches such as MotionMapper^27^, B-SOiD^28^, and keypoint-MoSeq^29^ are powerful for discovering pose-dynamic structure without predefined labels. Recent rodent studies have begun to combine behavioral phenotyping with interpretable multivariate and machine learning frameworks, including PCA-derived severity axes^62^, random-forest Gini-based variable importance^63^, SVM-based SHAP analysis^64^, and random-forest-based SHAP analyses^65,66^ to identify genotype-, disease-, and drug-associated behavioral signatures. Within this landscape, Autobehaver extends explainable, group-discriminative phenotyping to *Drosophila* by preserving semantic interpretability at the frame level and converting behavior predictions into high-dimensional per-animal profiles. These profiles support pairwise group discrimination, continuous placement of intermediate phenotypes, and SHAP/FSEA-based prioritization of the individual features and feature groups that define group-specific behavioral signatures. Thus, Autobehaver connects labeled behavior sequences to interpretable comparative phenotyping, providing a framework for discovering candidate behavioral modules in genetic, circuit, aging, and future suppressor or drug screens.

## Methods

### Imaging Platform and Pi Housing

We designed and built a prototype Imaging Platform based on low-cost, scalable recording hardware. A full parts list is found in Supplemental Table 20. Individual recording units consisted of an Arducam MINI High Quality 12.3 MP 1/2.3” 477P Camera with 12 mm Low Distortion M12 lenses (Arducam Co., Ltd., Nanjing, Jiangsu, China), connected to a Raspberry Pi 5 unit (RPi; Raspberry Pi Ltd, Cambridge, UK). Each recording unit was connected to a Catalyst WS-C2960X-24TS-L network switch (Cisco, San Jose, CA), allowing centralized control from a Precision 7510 laptop (Dell, Round Rock, TX). The cameras were mounted facing downward inside the Imaging Platform.

### Live Recordings

Plates with small chambers for flies were 3D printed using the Bambu Lab X1E with Ivory White Matte PLA (Bambu Lab, Shenzhen, China). The plate was designed such that each camera observed a bank of six chambers. Each chamber is 2.5 mm deep and 9 mm in diameter with slightly tapered walls to avoid camera occlusion while recording. Flies were anesthetized on ice by burying a vial in ice for 5 minutes and then placed into chambers using a staggered pattern of alternating genotypes and sexes. For each trial of an experiment, the pattern was shifted so that each camera saw each genotype in a different chamber position. After loading, the chambers were sealed with a cover plate of transparent 1/16-inch acrylic (Piedmont Plastics, Inc., Charlotte, NC, USA), and the plate was placed inside the imaging box for at least 30 minutes prior to recording so that the flies could recover and acclimate. All genotypes were collected, aged, and handled in parallel to ensure identical treatment across groups.

Recordings were coordinated by the central laptop computer, which sent commands to all RPis simultaneously via a custom Python script. Each RPi recorded video at 2016 × 1520 pixel resolution and 40 frames per second (FPS) for between 26-30 minutes. Each video consisted of a full camera view with six chambers. After recording, video files were transferred to a network drive for storage, and copied onto the high-performance computing cluster (Biowulf) for data processing. The first processing step used a custom Python script to detect chambers with a Hough circle transform and crop each chamber to a 600 × 600-pixel movie using OpenCV (OpenCV Team, version 4.12).

#### Drosophila melanogaster

*Drosophila melanogaster* strains were raised on Bloomington Recipe Fly food from LabExpress (Ann Arbor, MI). All experimental fly crosses were performed at 25°C, either in vials with 8-12 females and at least half as many males, or in bottles with 20-25 females and at least half as many males. Flies were collected at 0-2 days old, kept in vials with 8 males and 8 females, and aged for 2, 15, 30, or 60 days. Details of all fly strains used are in Supplemental Table 21.

### DeepLabCut *Drosophila* Keypoint-tracking Model

We trained a custom DeepLabCut^25^ (DLC) model to predict keypoints (x, y, and likelihood values) for 30 body parts for every frame of a given movie. Training data for DLC included 2968 annotated images of 571 flies, including males and females of six visually distinct phenotypes (Supplemental Table 1). The 30 keypoints tracked included multiple points across the head, thorax, abdomen, legs, and wings (Supplemental Figure 1). In order to calculate the animal centroid and angle, we included both a Front and Back keypoint that were always labeled in our training data regardless of whether they were visible or occluded. We trained our DLC model using DeepLabCut 2.3.9 with a ResNet-50 backbone for 1,000,000 iterations, resulting in a median test error of 3.62 pixels or 0.062 mm (Supplemental Table 2).

### Data Pre-processing

Each cropped movie was processed by our DLC model^25^ to generate a matrix where each row represents a movie frame and each column represents an x, y, or likelihood value for a given body part. Raw DLC files were pre-processed to prepare for neural network input, including transforming keypoints into an egocentric reference frame and extensive feature engineering for a total of 820 features. Z-score normalization was applied to each feature column. See Supplemental Methods for full details and all formulae.

### Neural Network Architecture and Training

We developed a Transformer encoder-based neural network architecture that uses pre-processed features to predict Behavior and Orientation labels (Supplemental Video 1). The architecture was constructed using PyTorch^67^ and the code was written in Python (Python Software Foundation). The input to the model is an odd number of rows from the pre-processed dataset, where the middle row of the input sequence is the frame of interest, and the rows before and after are the past and future temporal context, respectively. The input sequence is passed through a linear input layer that transforms the input rows into a common vector space to become “frame tokens”. We optionally append a learned class token (CLS) to the sequence. We added either a sinusoidal or learned positional encoding to each frame token’s embedding. These tokens are then passed through one or more Transformer encoders. After Transformer encoder processing, either the middle frame token or the CLS token is isolated as the latent representation of the input sequence.

We performed supervised training, where the objective is to use the latent representation to predict Behavior and Orientation labels for the middle frame. Augmentations were applied to each feature in the input sequence during batch construction. First, Gaussian noise was added to all input features. Second, masking was applied independently to each input feature, such that each feature had some probability of being set to zero. Each augmented input was passed through the encoder network to generate the latent representation, which was then passed through two multi-layer perceptron prediction heads to yield probability distributions for the pre-defined Behavior and Orientation labels (Supplemental Figure 1). Cross-entropy loss was calculated by comparing the output probabilities with ground-truth labels. Since our labelled dataset contained a different number of examples for each BxO combination, we calculated class weights that were inversely proportional to the frequency of each BxO class in the dataset. These class weights were used to adjust model updates in order to correct for over/under-represented classes. Initial model weights were initialized as follows: Xavier uniform random initialization was used for input linear projection, classifier-head fully connected layers or multi-layer perceptron layers, Transformer feedforward, and attention projection weights; LayerNorm weights were initialized to 1; learned positional embeddings and CLS tokens were initialized from a small normal distribution with mean 0 and standard deviation 0.02; and all biases were initialized to 0. The network was trained with a learning rate that followed a single-epoch linear warmup to a user-defined maximum learning rate (typically 1e-4) followed by a 300-epoch cosine decay schedule (final learning rate 1e-6) using the AdamW optimizer.

Our dataset contained 385,933 total frames, corresponding to roughly 2.7 hours of video, with corresponding Behavior and Orientation labels. Video clips were manually annotated using a custom script with a graphical user interface that displays full-length videos and supports bout-based annotation. The custom script supports two annotation modes. In the first mode, the user selects a segment of the full-length video and annotates the start and end frames of sequential Behavior and Orientation bouts within that segment. In the second mode, the script also incorporates inference results from a previously trained model to highlight clips containing either rare bouts or low-confidence predictions, thereby allowing targeted annotation. A standard training-validation split was generated at the level of video clips and was used for all architecture tuning and dataset ablation experiments. We generated 100 random clip-level splits targeting approximately 80% of labeled frames in the training set and 20% in the validation set, ranked these candidate splits according to the balance of combined BxO labels between the two partitions, and selected a top-ranked split that retained at least one example of each combined BxO label in the validation set.

Architecture tuning and dataset ablation experiments were performed before training final deployment models using the standard training-validation split described above and a fixed set of random seeds (Supplemental Figures 2 and 3). For each run, the best-performing epoch was selected based on validation macro F1 for Behavior prediction. After the final architecture was chosen, final deployment models were retrained on the full labeled dataset including all calculated features and without a held-out validation set in order to maximize data usage, retaining the checkpoint from the final epoch. Architectural parameters for the three final models are listed in Supplemental Table 12.

We performed inference using an ensemble of three trained models. For each model, we performed test-time augmentation (Gaussian noise level 0.01 applied to all input features) and averaged the logits from three forward passes of each of the three models. Softmax was applied to the averaged logits to obtain probability distributions over the Behavior and Orientation labels. The resulting label sequences were then post-processed with a minimum bout length of four frames. Specifically, non-Jump bouts shorter than four frames were reassigned based on their flanking labels: if the same label occurred on both sides of the short bout, the intervening frames were set to that label; if the flanking labels differed, the short bout was reassigned to the flanking label with the greater summed predicted probability across those frames. Jump bouts were exempt from smoothing as they occur over 1-3-frame intervals.

### Post-inference Feature Engineering

After generating BxO predictions for an experimental dataset, we performed extensive post-inference feature engineering to construct a high-dimensional per-animal feature matrix from predicted bout structure and keypoint-derived measurements. This matrix integrates bout summary, time budget, body measurement, postural, kinematic, transition, sleep, and spatial feature categories, including both bout-based and context-level summaries. Sparse bout-dependent features were masked when insufficient data were available for the corresponding context. See Supplemental Methods for full details and formulae, and Supplemental Table 13 for the full list of features.

### XGBoost Training and Feature-Importance Analysis

We calculated the full behavioral feature matrix for every individual within an experiment and performed a feature-importance analysis. The first step in this analysis was to define a normalization reference group. Then, for each feature, we calculated the mean and standard deviation of the normalization reference group, and used those parameters to perform z-score normalization for every feature of every animal in the experiment. Because some individuals lacked certain features, we filtered to remove any feature where fewer than 25% of individuals in either reference group had a given feature.

To quantify group-discriminative behavioral features, we trained ensembles of binary XGBoost classifiers^49^ under a random subsampling design with multiple rounds. In each round, we partitioned the full feature set into random subsets of 100 features, such that each feature was assigned to one subset in that round whenever possible. When the total number of features was not divisible by 100, the final subset was padded to the target size by randomly resampling features from the full feature list. For each round we also generated a fresh five-fold, group-stratified cross-validation split at the level of individual animals. For every feature subset in a given round, we trained one model per fold, fitting on four folds and evaluating on the held-out fold. This procedure was repeated for 100 rounds so that each feature was sampled repeatedly across the ensemble. In parallel, we repeated the same procedure using permuted training-fold labels within sex strata while leaving held-out test labels unchanged, thereby generating a negative-control null distribution. For each real feature subset, we pooled held-out predictions across its five fold-models and summarized out-of-fold performance as the normalized improvement in log loss relative to a trivial baseline classifier that ignored all features and predicted the training-fold class prevalence for every animal. We then used the corresponding distribution from the permuted runs to assign each subset a non-negative performance weight, so that subsets exceeding the permuted null contributed more strongly to the final ensemble:

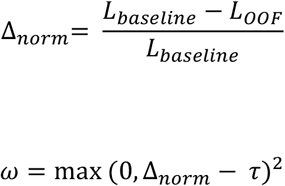

Where L_OOF_ is the pooled out-of-fold log loss for a feature subset, L_baseline_ is the fold-aware constant-baseline log loss, and τ is the permutation-derived null threshold. In practice, subsets that outperformed the permutation-derived null received greater weight in both prediction aggregation and SHAP summarization. Final ensemble predictions were then obtained by reloading the saved fold-models, aggregating their raw margins in logit space using these subset-level weights, and converting the aggregated margins to probabilities with a sigmoid transform.

For analyses of intermediate phenotypes, we linearly rescaled each individual’s final ensemble class prediction using the mean out-of-fold ensemble predictions of the two reference groups. This produced a Phenotype Score (PS) in which the mean predictions of the two reference groups defined the endpoints of the scale, and intermediate values reflected relative position along the classifier-defined discriminative axis. Behavioral feature vectors from non-reference groups were passed through the same weighted ensemble and rescaled using the same reference-group calibration parameters. Because these calibration parameters were defined separately within each experiment, PS values were interpreted only within a given dataset.

To quantify feature importance within the classifier ensembles, we computed SHapley Additive exPlanations (SHAP) values for each trained XGBoost model ^52^. For tree-based models, SHAP provides an additive decomposition of the model output into an expected value plus per-feature contributions for each individual animal. Positive SHAP values indicate that a feature pushes the prediction toward the positive class and negative values indicate the opposite. To ensure consistency with the weighted ensemble prediction scheme, we reloaded the saved fold-specific models and recomputed SHAP values using the same subset-level performance weights used during prediction aggregation. For training individuals, out-of-fold SHAP values were constructed by retaining only the held-out fold contribution for each animal; for non-training individuals, SHAP values were obtained by averaging contributions across folds within each subset. Because each XGBoost model was trained on only a random subset of features, SHAP values were defined only for features present in a given model. Accordingly, for each feature we aggregated SHAP values only across the subset-models in which that feature was included, using feature-specific weighted denominators so that features were compared only across the weighted subsets in which they were observed.

We summarized importance in two complementary ways. First: group-specific mean SHAP, defined as the mean signed out-of-fold SHAP value within each training group. Second: global importance, defined from the mean absolute weighted out-of-fold SHAP value across both training groups after subtraction of the permutation-derived SHAP baseline and clipping at zero. For non-training individuals, averaged SHAP values were computed across the same weighted ensemble, enabling direct comparison of feature-level deviations with the training-group profiles.

As an independent, unsupervised analysis, we performed principal component analysis (PCA) on the same normalized behavioral feature matrix used for classifier training. We then defined a reference-group axis as the line connecting the centroids of the two reference groups in PCA space. Each individual was decomposed into an on-axis coordinate, given by its projection onto this line, and an off-axis value, given by its perpendicular distance from the line. Because both quantities were measured in the same PCA coordinate system, they were reported on the same unitless scale.

### Feature Set Enrichment Analysis

To identify higher-order structure in the set of important features, we performed Feature Set Enrichment Analysis (FSEA), adapted from gene set enrichment analysis^53^ (GSEA). First, we ranked all features by their SHAP-based global importance. We then tested whether predefined feature sets were over-represented near the top of this ranked list using the pre-ranked GSEA procedure implemented in GSEApy (gseapy.prerank). Feature sets were defined *a priori* by grouping related features into interpretable categories (e.g., time budgets, bout statistics, transitions, kinematics, posture, spatial, and sleep) within specific Behavior and/or BxO contexts (Supplemental Table 14).

For each feature set, FSEA computes an enrichment score using a weighted running-sum statistic over the ranked feature list and estimates statistical significance via permutation testing (100,000 permutations) as implemented in GSEApy, reporting normalized enrichment scores and multiple-testing-corrected FDR q-values. Feature sets with FDR ≤ 0.1 were considered significantly enriched.

### Comparison of Subsampling and Full-feature XGBoost Approaches

Our primary feature-importance framework used repeated random feature subsampling, such that each XGBoost model in the ensemble was trained on only 100 features at a time. To test whether this design provided a practical benefits for feature importance analysis, we compared it with a standard full-feature implementation consisting of ensembles of 100 XGBoost models trained on the complete behavioral feature matrix without feature subsampling. For each dataset (*UBA5* and *spd-2*), both methods were repeated three times with independent random seeds.

We first compared classification performance using area under the curve (AUC) and F1 score. To assess how concentrated feature importance was along the ranked feature list, we plotted the cumulative fraction of total global importance captured as features were added from highest to lowest rank. To evaluate reproducibility of the ranked feature lists across runs, we used two complementary metrics: Jaccard overlap of the top-k feature sets and Spearman rank correlation on the union of the top-k feature sets. Finally, we compared the recovery of enriched feature groups by FSEA (reported as footnotes in Supplemental Tables 17 and 19).

## Supporting information

All Supplemental Tables

10 animal poses

example inference

## Acknowledgements

We thank Hong Xu and Brian Galletta for feedback on the project. We thank Ibraheem Farooq, Marcial Garmendia-Cedillos, Ghadi Salem, Brian Oliver, and Jason Tennessen for feedback and discussions related to design and operation of the video recording platform, and application of keypoints to behavioral recordings. We thank Sarah Hooper, Ed Giniger, and Susan Harbison for feedback on a final draft of the manuscript. This work utilized the computational resources of the NIH HPC Biowulf cluster (https://hpc.nih.gov). This work was supported by the Division of Intramural Research at the National Heart, Lung, and Blood Institute (ZIAHL006126 to NMR).

## Disclaimer

The contributions of the NIH author(s) are considered Works of the United States Government. The findings and conclusions presented in this paper are those of the author(s) and do not necessarily reflect the views of the NIH or the U.S. Department of Health and Human Services.

## Data availability statement

All relevant data can be found within the article and its supplementary information. All raw data will be made available under the Digital object identifier (DOI)10.25444/nhlbi.32180805

## Code availability statement

All code will be made available through the Nature Communication submission system. Following review, all code will be made available on GitHub.

---

**Supplemental Figure 1:**
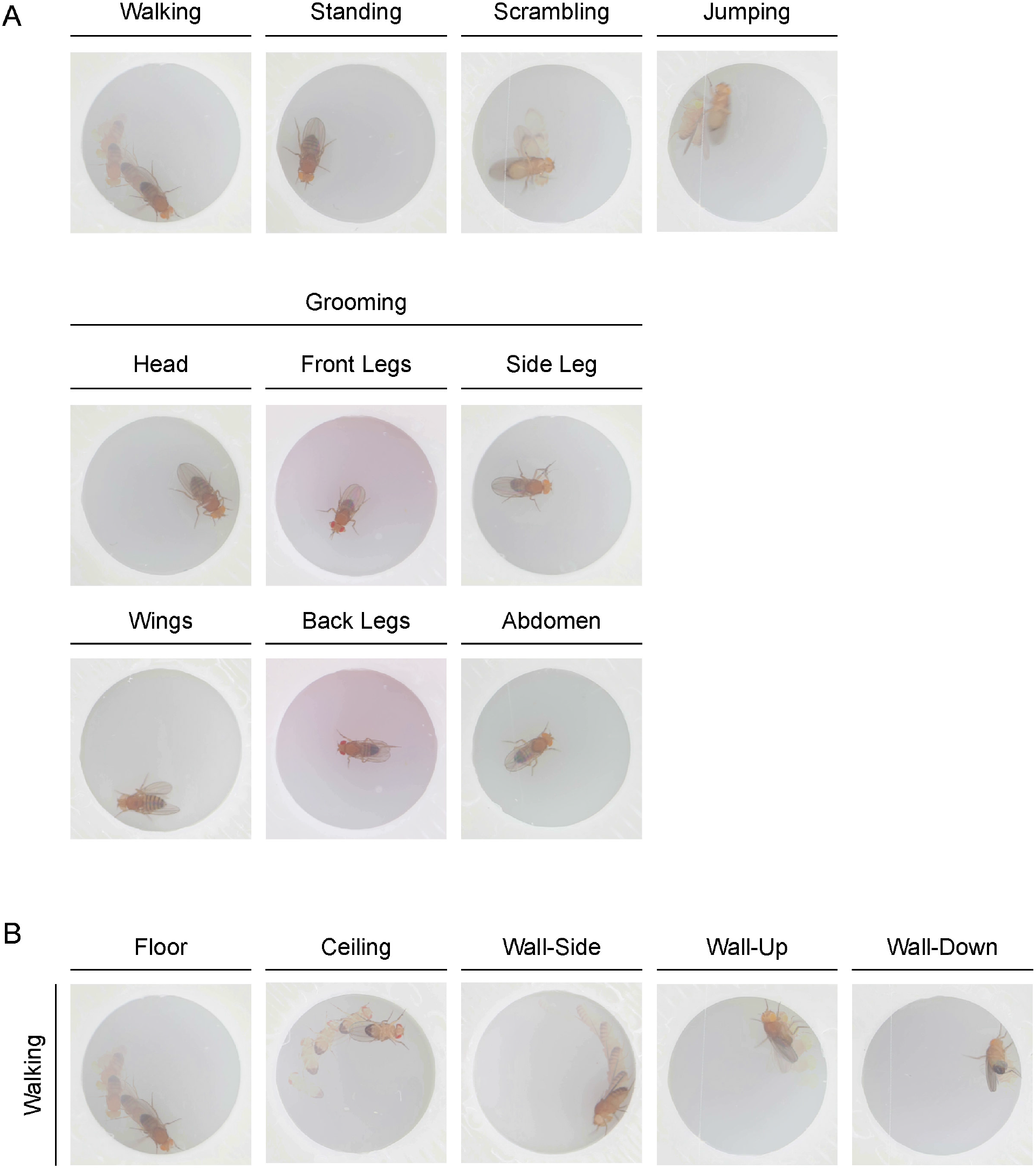
Behaviors and Orientations The 10 annotated Behaviors (A) included Walking, Standing, Scrambling, Jumping, and six types of Grooming: Head, Front Legs, Side Leg, Wings, Back Legs, and Abdomen. The five annotated Orientations (B) are shown in combination with Walking Behavior. Dynamic poses are shown in stroboscopic view to depict movement.

**Supplemental Figure 2:**
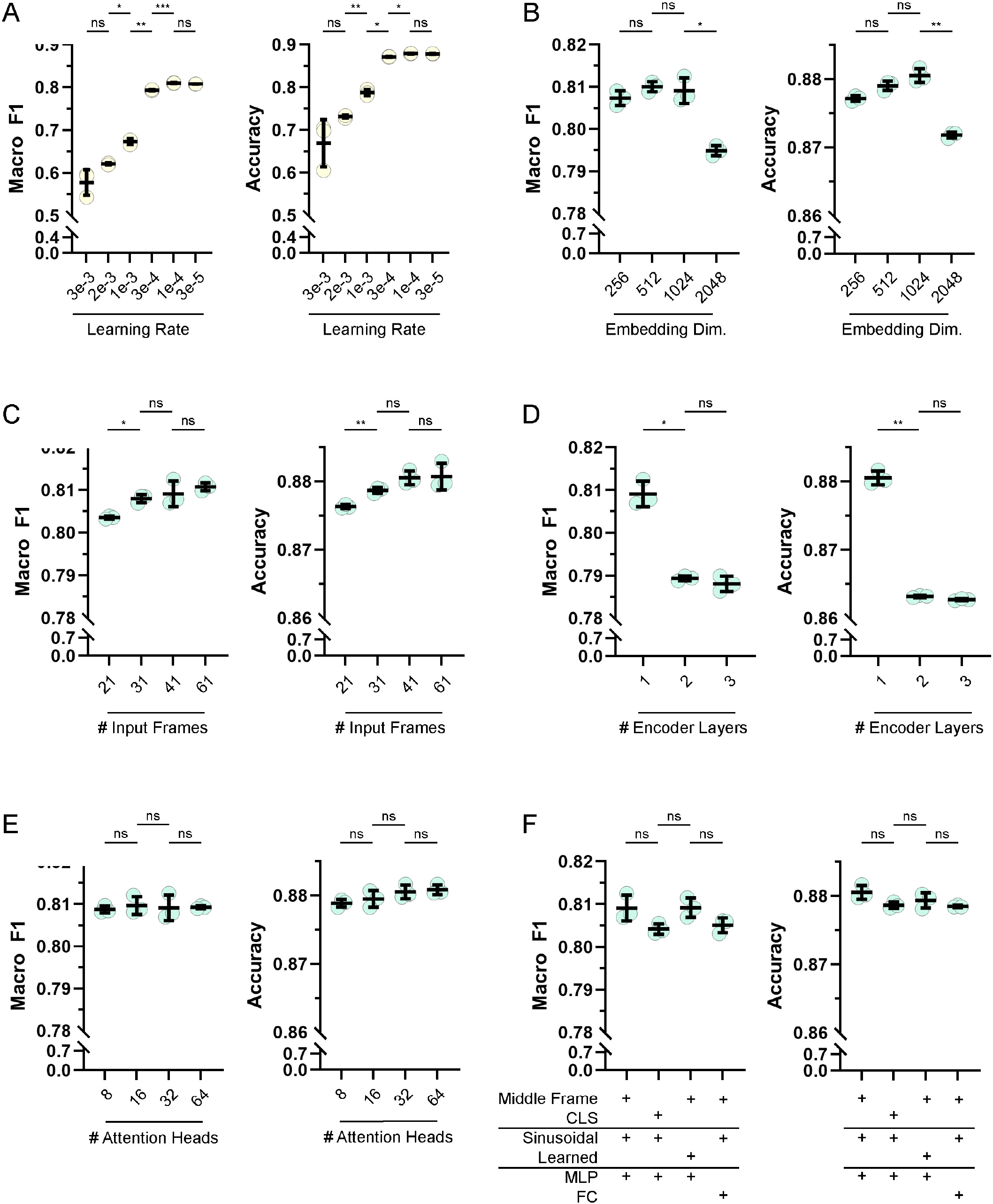
Hyperparameter and Architecture Tuning Results Architecture tuning experiments were performed using a dataset containing all features, a predefined training-validation split, predefined random seeds, and a common set of architecture parameters with a single change for each experiment. Results are reported as Validation Macro F1 (A, C, E, G, I, K) and Validation Accuracy (B, D, F, H, J, L) for three replicates, with column labels indicating the altered parameter(s). First, base learning rates were tested (A, B) using common parameters of 41 input frames, one encoder layer, an embedding dimension of 512, 32 attention heads, sinusoidal positional encodings, a middle-frame representation, multi-layer perceptron (MLP) prediction heads, and a 300-epoch cosine decay schedule with a final learning rate of 0.000001. Second, embedding dimensions (C, D) were tested using the same common parameters and base learning rate 0.0001. Third, the number of input frames (E, F), encoder layers (G, H), and attention heads (I, J) were tested using the same common parameters, except embedding dimension which was set to 1024. We also tested CLS token instead of middle frame token representation, learned instead of sinusoidal positional encodings, and fully connected (FC) instead of MLP prediction heads (K, L). Statistical comparisons were made using Welch’s ANOVA with Dunnett’s T3 post-hoc test for all-to-all pairwise comparisons. *** p < 0.001; ** p < 0.01; * p < 0.05; ns, not significant. Only selected statistical results are shown for simplicity.

**Supplemental Figure 3:**
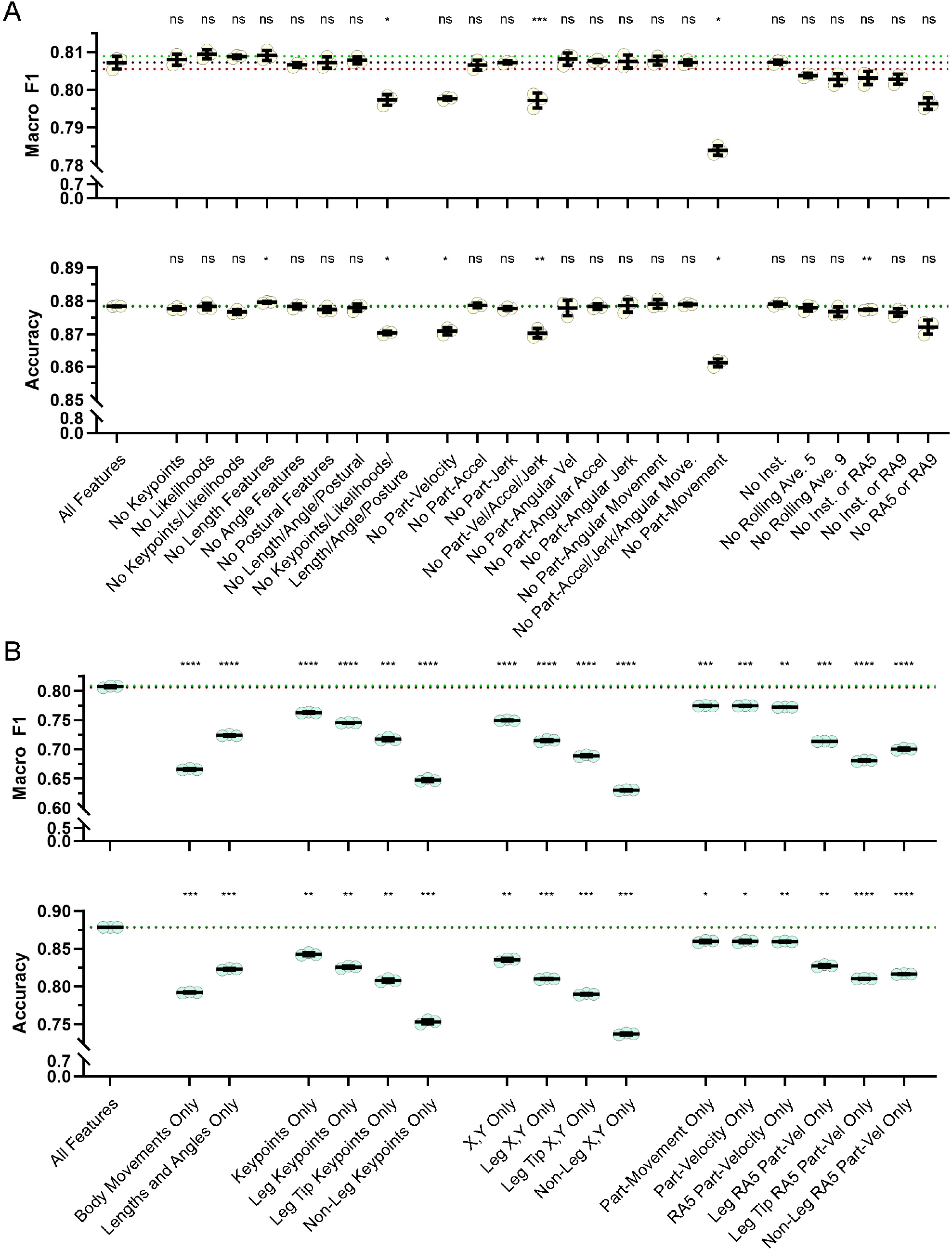
Dataset Ablation Experiments Dataset ablation experiments were performed using a predefined training-validation split, predefined random seeds, and a common set of architecture parameters: 41 input frames, one encoder layer, an embedding dimension of 1024, 32 attention heads, sinusoidal positional encodings, middle-frame representation, multi-layer perceptron prediction heads, and a 300-epoch cosine decay schedule with base learning rate 0.0001 and final learning rate of 0.000001. Results are reported as both validation Macro F1 (top) and Accuracy (bottom) for each of three replicates, with the mean and upper and lower standard deviation bounds of the “All Features” model shown as black, green, and red dotted lines, respectively. Experiments were performed by either dropping feature subsets (A) or by training on only one feature subset (B). Statistical comparisons were made using Welch’s ANOVA with Dunnett’s T3 post-hoc test for pairwise comparisons against “All Features”. **** p < 0.0001; *** p < 0.001; ** p < 0.01; * p < 0.05; ns, not significant.

**Supplemental Figure 4:**
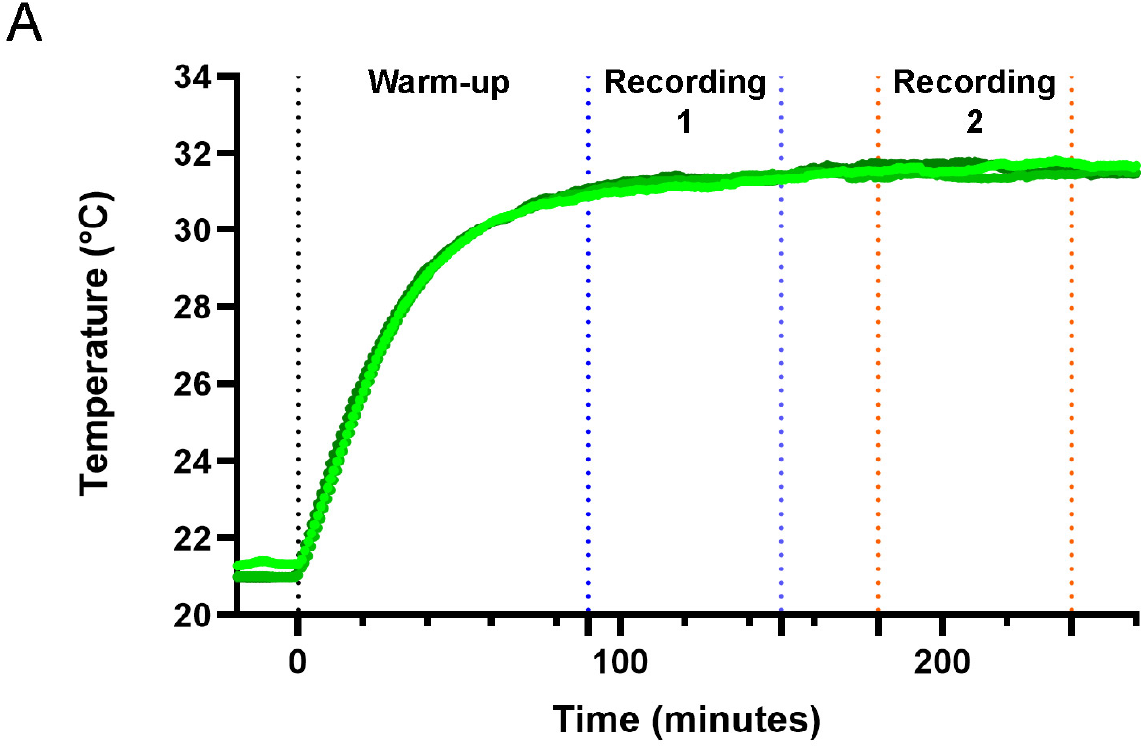
Temperature During Recording Temperature recordings from inside the imaging box showed temperature eventually plateaued at 31.8°C (exponential one-phase association regression, R^2^ 0.965). The first recording could be initiated as early as 90 minutes, at which point temperature rose from 31°C to 31.4°C over the recording period. The second recording could be initiated as early as 180 minutes, when temperature was stable at 31.6°C.

**Supplemental Figure 5:**
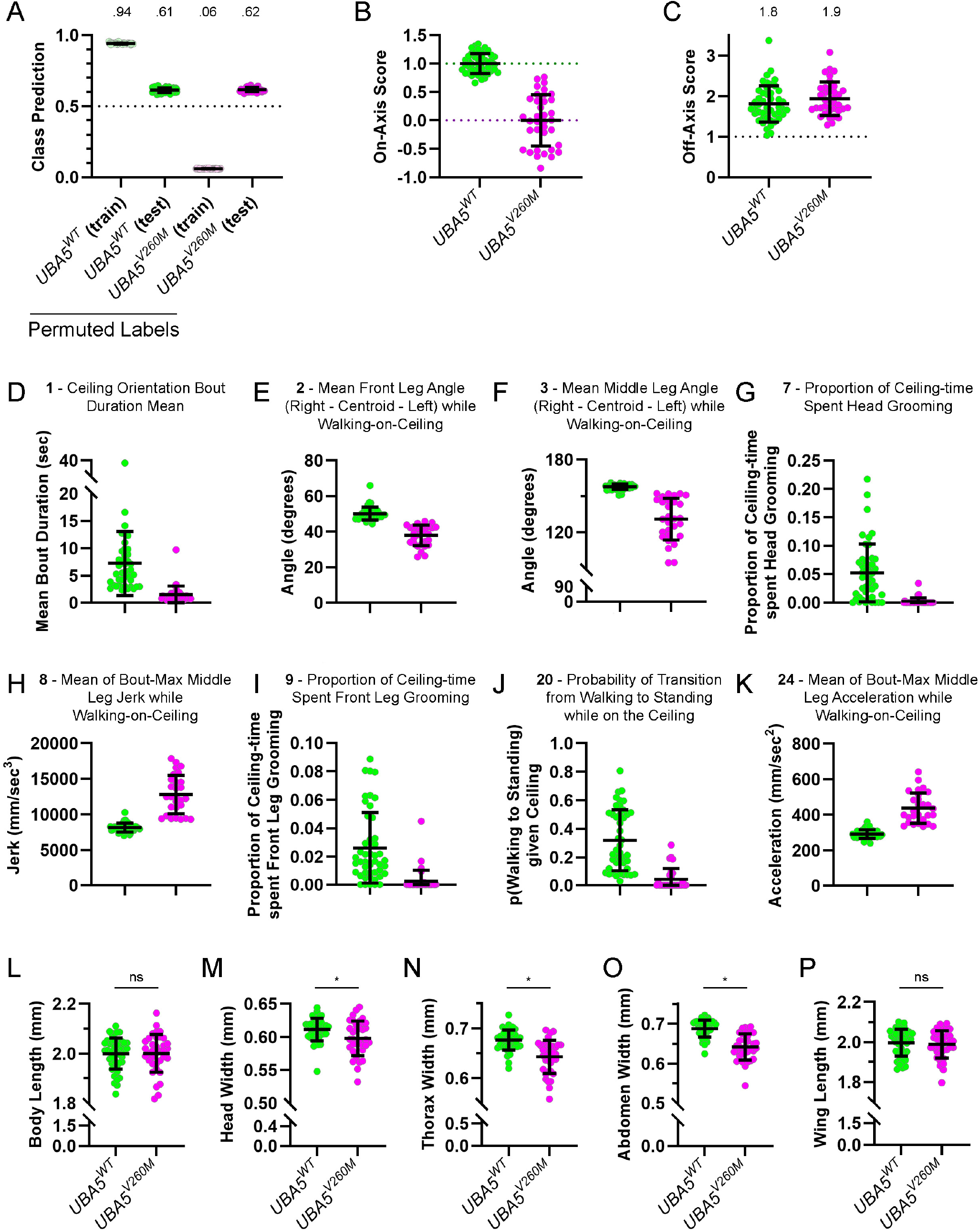
*UBA5* Supporting Information Training a classifier ensemble on *UBA5* individuals with permuted labels (A) resulted in no discriminative power (average class prediction for non-reference individuals = 0.62). The full behavioral feature matrix was projected into principal component space to define an axis separating the centroids of *UBA5*^*WT*^ and *UBA5*^*V260M*^ individuals, resulting in on-axis (B) and off-axis (C) components for each individual, both reported in the same unitless scale. On-axis values indicate each fly’s position along the line connecting the average positions of the two reference groups in PCA space; off-axis values indicate each fly’s distance from that line. Raw feature plots for selected top-ranked Ceiling Orientation-related features are shown, with the number preceding each feature name indicating its rank in the global-importance list: ceiling bout duration mean (D); mean angle formed between the front right tarsus, centroid, and front left tarsus while Walking on the Ceiling (E); mean angle formed between the middle right tarsus, centroid, and middle left tarsus while Walking on the Ceiling (F); proportion of total Ceiling time spent Head Grooming (G); mean of bout-specific maximum middle leg jerk while Walking on the Ceiling (H); proportion of total Ceiling time spent Front Leg Grooming (I); probability of transitioning from Walking to Standing while on the Ceiling (J); and mean of bout-specific maximum middle leg acceleration while Walking on the Ceiling (K). Standard measurements of *UBA5*^*WT*^ and *UBA5*^*V260M*^ individuals are shown for body length (L), head width (M), thorax width (N), abdomen width (O), and wing length (P). Statistical significance was determined by Welch’s t-test. * p < 0.05; ns, not significant.

**Supplemental Figure 6:**
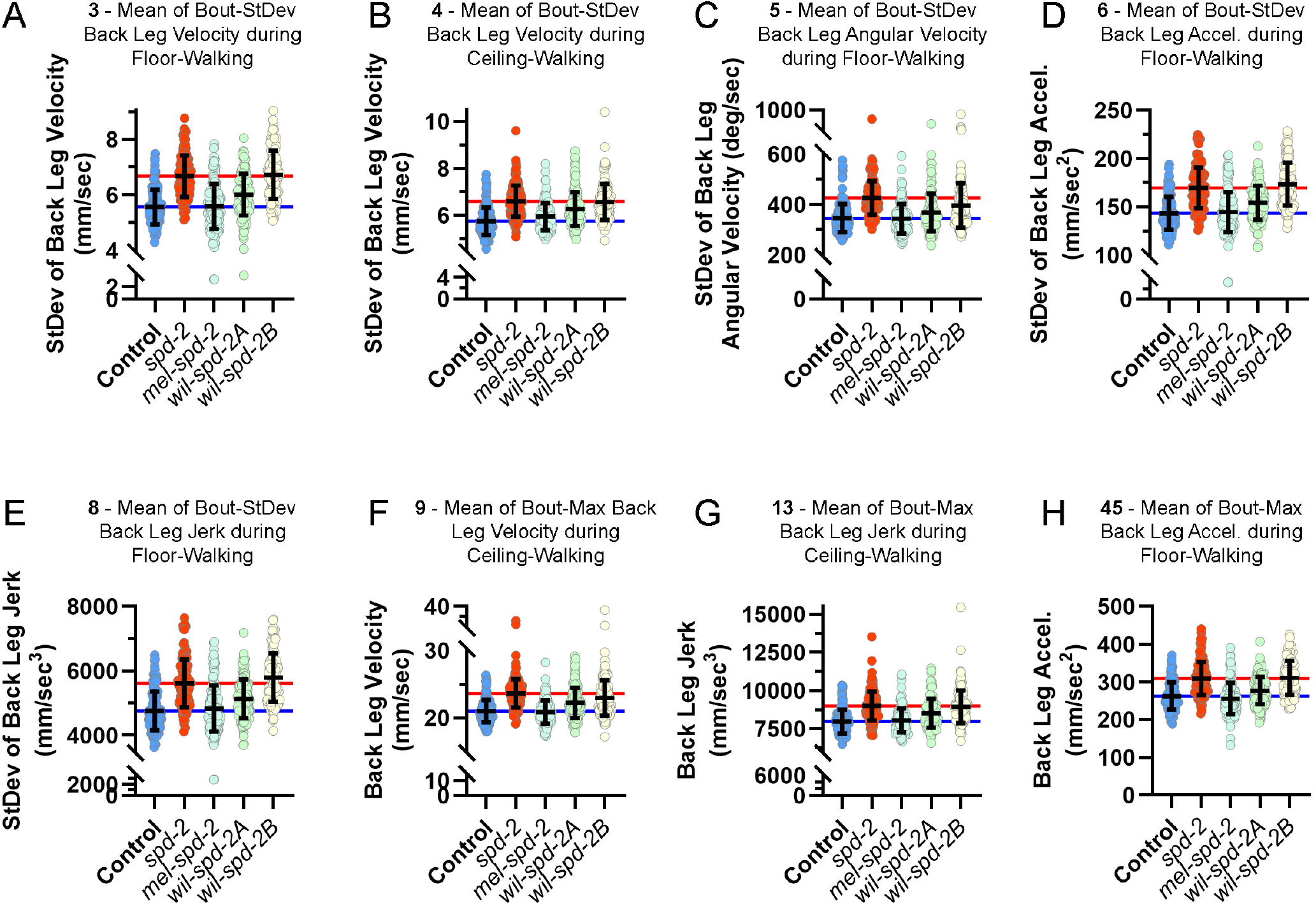
*Spd-2* Supporting Information Raw feature plots for selected top-ranked features from the feature group describing back leg kinematics during Walking bouts (A-H). The number preceding each feature name indicates its rank in the global importance-ranked list. Horizontal colored lines indicate the mean values for control (blue) and *spd-2* (red).

**Supplemental Figure 7:**
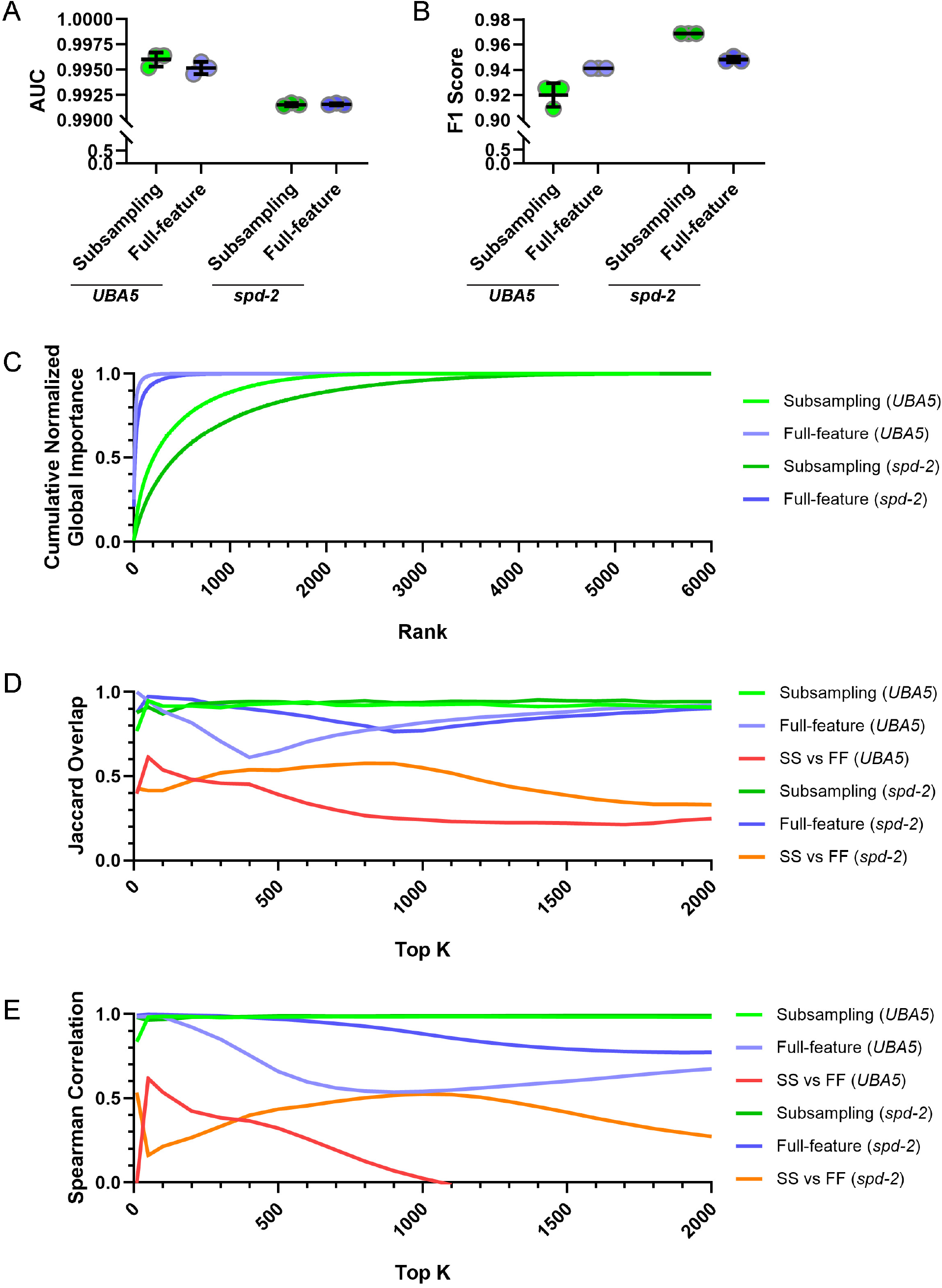
Comparison of Subsampling and Full-feature XGBoost Feature-Importance Strategies Comparison of the subsampling XGBoost and full-feature XGBoost approaches on the *UBA5* and *spd-2* datasets. Classification performance across three independent runs is shown as AUC (A) and F1 score (B). Cumulative normalized global importance (C) shows that full-feature XGBoost concentrates importance in a small number of top-ranked features, whereas the subsampling approach distributes importance more broadly across the ranked list. Jaccard overlap among the top-k globally important features across the three independent runs is shown for the *UBA5* and *spd-2* datasets (D), along with overlap between the subsampling and full-feature approaches within each dataset. Spearman rank correlation on the union of the top-k feature sets is shown for the same comparisons (E). In D and E, subsampling is shown in green, full-feature XGBoost in purple, and between-method comparisons in red/orange, with lighter colors indicating the *UBA5* dataset and darker colors indicating the *spd-2* dataset. SS, subsampling XGBoost; FF, full-feature XGBoost.

## Additional Supplemental Material

### Supplemental Methods File

**Supplemental Tables 1-21**

**Video File 1:** Examples of the 10 Behavior labels, shown as Floor, Ceiling, and Wall-Side Orientations.

**Video File 2:** Example inference of *Drosophila* Behavior and Orientation labels.

## Supplemental Methods

### Data Pre-processing

Each cropped movie was processed by DeepLabCut, converting the movie into a spreadsheet where each row represents a movie frame and each column represents an x y, or likelihood value for a given body part. Raw DLC files were further pre-processed to prepare for neural network input. First, centroid (*C*_*x*_ and *C*_*y*_) position was calculated using the Front (*F*) and Back (*B*) keypoints:

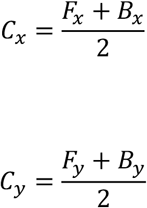

Next, body angle (*θ*_degrees_) was calculated using the *F* and *B* keypoints:

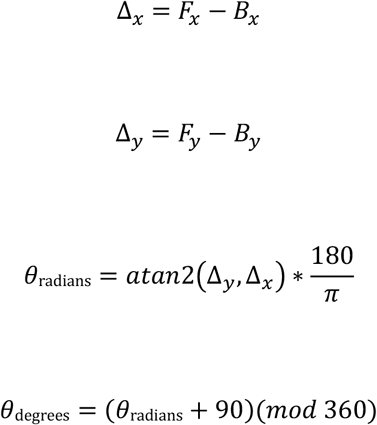

We performed a series of corrections on the raw x y keypoints to separate the overall position and angle of the fly from the relationships between body parts. First, we subtracted the centroid coordinates *C*_*x*_ and *C*_*y*_ from all raw keypoints *P*_*x*_ and *P*_*y*_, pinning the centroid at coordinate (0, 0):

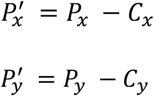

Second, we rotated all parts around the center of the plane based on the current angle such that the fly always points North:

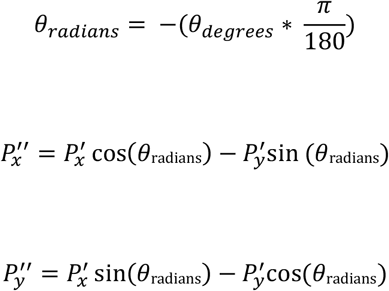

We engineered multiple additional features to include as input data. Overall animal movement features and part-specific movement features were calculated based on the average change from the previous frame to current frame and current frame to next frame, using raw centroid position or corrected part-specific position, respectively. First, a 2D movement vector (*M*_*t*_) was calculated for each frame t using the centroid positions (*C*) from the preceding (t − 1) and subsequent (t + 1) frames:

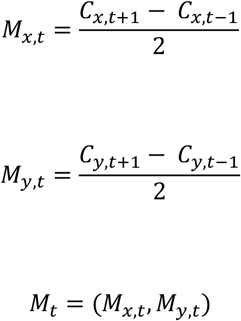

This instantaneous movement vector was then used to create three separate features. Velocity (magnitude; *v*_*mag,t*_) represents the overall speed of the animal or part, regardless of direction:

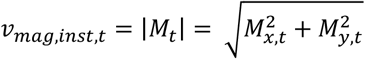

Forward velocity (*v*_*fwd,t*_) and strafe velocity (*v*_*strafe,t*_) represent the component of the animal’s movement that is parallel and perpendicular to the body axis, respectively, and were calculated for the whole body movement. First, the body axis vector *A*_*t*_ is defined using the Back (*B*) and Front (*F*) keypoints:

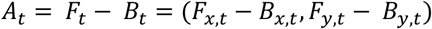

Additionally, we defined a pair of perpendicular axes using Left/Right (*A*_*LR*_) and Top/Bottom (*A*_*TB*_) keypoints. For each frame, the perpendicular axis used is based on which one has the higher summed keypoint likelihood:

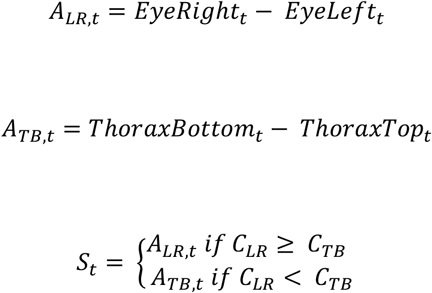

These vectors are normalized to create unit vectors (*Â*_*t*_ and *Ŝ*_*t*_) that point in the forward and strafe directions, respectively:

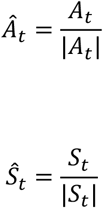

Finally the forward velocity is calculated by projecting the *M*_*t*_ onto the normalized vectors *Â*_*t*_ and *Ŝ*_*t*_:

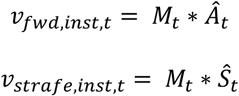

Acceleration was calculated based on the change in *v*_*inst,t*_ (using *v*_*mag,t*_, *v*_*fwd,t*_, and *v*_*strafe,t*_ to calculate *a*_*mag,t*_, *a*_*fwd,t*_, and *a*_*strafe,t*_, respectively) using the formula:

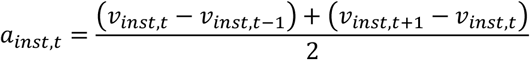

Jerk was similarly calculated for the animal centroid based on the change in *a*_*inst,t*_ (using *a*_*mag,t*_, *a*_*fwd,t*_, and *a*_*strafe,t*_ to calculate *j*_*mag,t*_, *j*_*fwd,t*_, and *j*_*strafe,t*_,, respectively) using the formula:

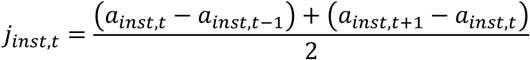

Next, angular velocity was calculated as the change in angle between a “moving” keypoint as it rotates around a “base” keypoint. Whole body angular velocity was calculated using the Front (moving) and Back (base) keypoints, and part-specific angular velocities were calculated based on pairs listed in Supplemental Table 3, using the formula:

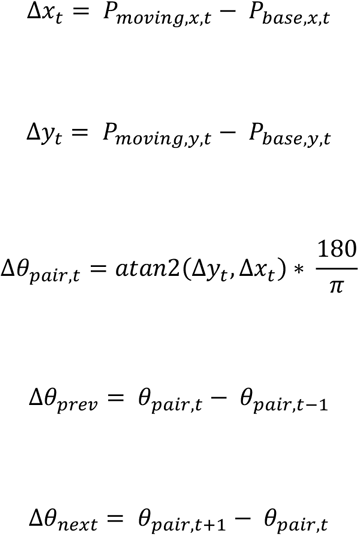

Both Δ*θ* values are normalized to the range [-180, 180], and used to calculate part-specific angular velocity *ω*_*pair,t*_:

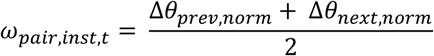

Angular acceleration (*α*_*inst,t*_) was calculated based on the change in *ω*_*pair,t*_ using the formula:

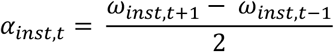

Angular jerk (*j*_*ang,pair,t*_) was calculated based on the change in *α*_*inst,t*_ using the formula:

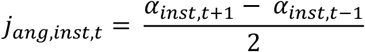

Path curvature was calculated as the ratio of the animal’s angular velocity to its forward velocity:

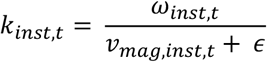

Rolling averages for all velocity, acceleration, jerk, angular velocity, angular acceleration, angular jerk, and curvature were calculated by averaging the instantaneous values from five or nine consecutive frames, centered on the current frame. We also calculated standard deviation features based on the instantaneous values across five- and nine-frame windows for all velocity and angular velocity features.

To help the model learn a fly’s orientation, we calculated a series of log likelihood ratios *λ*_*ratio*_ using the likelihood values of opposite body part pairs (*L*_1_ and *L*_2_; Supplemental Table 4), under the assumption that visible points will have high likelihood and hidden points will have low likelihood:

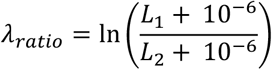

We calculated static length features as the distance between selected pairs of keypoints (Supplemental Table 5):

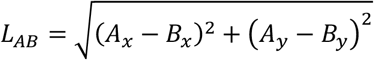

We calculated static angle features (*ψ*) as the angle formed by three keypoints (a base B and two limbs A and C; Supplemental Table 6):

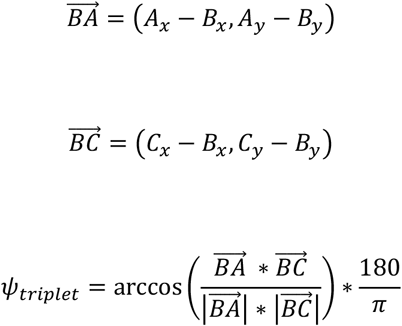

Finally, we calculated head orientation angle features which measure the orientation of the head, based on either the Left and Right eyes (R and L) or the Ocelli and Mouthparts (O and M) relative to the main body F-B axis. When the fly is standing in neutral position, these features should equal 90°; as the fly looks to the right or downwards, the features approach 0°, and as the fly looks to the left or upwards the features approach 180°:

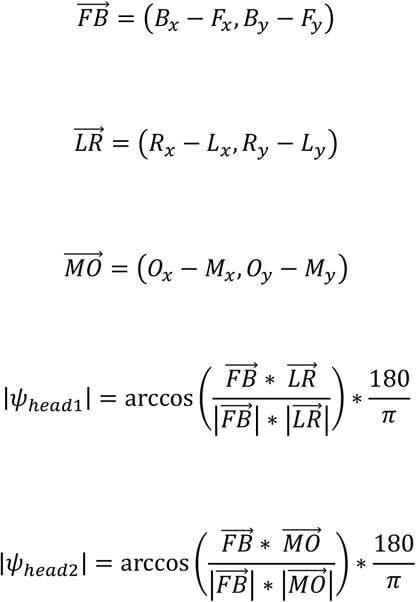

Area features were calculated as the area enclosed by specific sets of keypoints (Supplemental Table 7).

Asymmetry indexes were calculated to capture the left-right or top-bottom asymmetry of specific keypoint pairs (Supplemental Table 8) using the corrected keypoints:

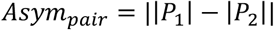

Altogether, the full pre-processed data contains 820 feature columns. Each column of data is z-score normalized across the entire dataset, such that mean is 0 and standard deviation is +/− 1. To optimize loading speed, the final normalized data files are saved in binary HDF5 format with 32-bit floating point precision.

### Post-inference Feature Engineering

After generating Behavior and Orientation (BxO) predictions for an experimental dataset, we performed extensive post-inference analysis to characterize behavioral differences between groups. The first step was to extract a high-dimensional feature matrix for each individual animal using a combination of bout and keypoint-based data. First, we extracted bouts of contiguous predicted labels: Behavior-only bouts, Orientation-only bouts, and BxO bouts. We then calculated high-level bout summary features. Bout frequency was calculated as the number of bouts divided by the total video time in seconds:

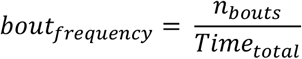

Mean bout duration was calculated:

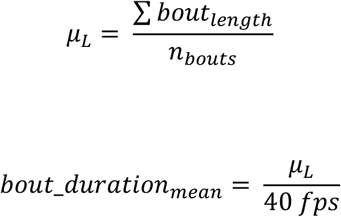

Bout duration coefficient of variation was calculated:

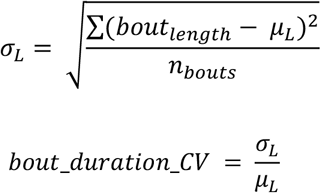

Initiation probability estimates the probability of starting a new bout when the animal is not already engaged in that bout:

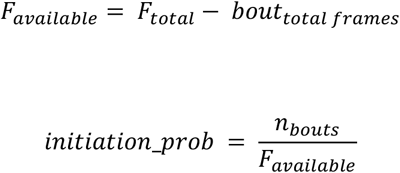

Inter-bout interval was calculated based on the time between the end of the previous bout and the start of the next bout of the same type.

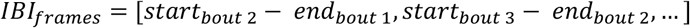

The mean and standard deviation of these intervals were then calculated and converted to seconds.

Next, time budgets were calculated for each bout type representing the total proportion of recorded time that the individual spent performing a given bout type:

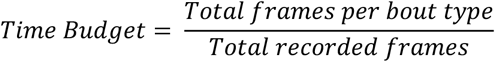

We also calculated conditional time budgets. *P*(*Behavior*|*Orient*) represents the proportion of time the individual spent performing a given Behavior relative to the total time spent in a specific Orientation. Conversely, *P*(*Orient*|*Behavior*) represents the proportion of time the individual spent in a given Orientation relative to the total time spent performing a specific Behavior:

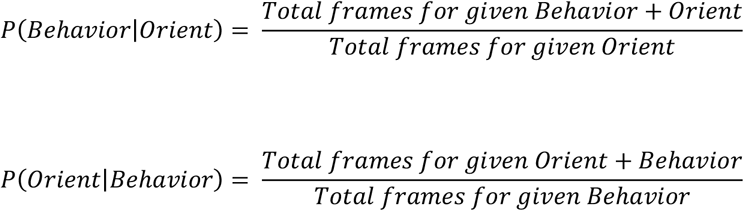

Standardized body-size features were calculated for each animal using Orientation-specific keypoint distances and body-area measurements (Supplemental Table 12). For each measurement, frames were first restricted to the required Orientation label and then searched using a Behavior hierarchy (Standing, Walking, Groom-FrontLegs, Groom-Head, Groom-SideLeg, Groom-Wings, Groom-Abdomen, Groom-BackLegs), stopping at the first Behavior yielding valid frames. From up to 100 valid frames per recording, we calculated the mean and standard deviation of each measurement. Measurement definitions included top-, bottom-, and side-view body length, head, thorax, and abdomen dimensions, left and right wing length, and body area. Recording-level values were averaged across recordings for each animal. Averaged body-size features were calculated by combining top- and bottom-view measurements, with side-view measurements used as fallbacks when top- and bottom-view data were unavailable. Normalized size features were obtained by dividing linear measurements, or the square root of body area, by the averaged thorax width. See Supplemental Table 12 for all measurements.

Postural features were calculated for each animal to characterize standing posture and body-part asymmetry. Postural features were extracted from Standing frames only, and were aggregated within each animal and Orientation by computing the mean and standard deviation across valid frames. For Floor- and Ceiling-oriented Standing frames, we retained only frames in which all required leg-tip and body-axis keypoints had likelihood > 0.1. We then calculated stance area as the polygonal area enclosed by the six leg tips, and calculated front-, mid-, and back-leg extension as the mean distance of each left-right leg pair from the body centroid:

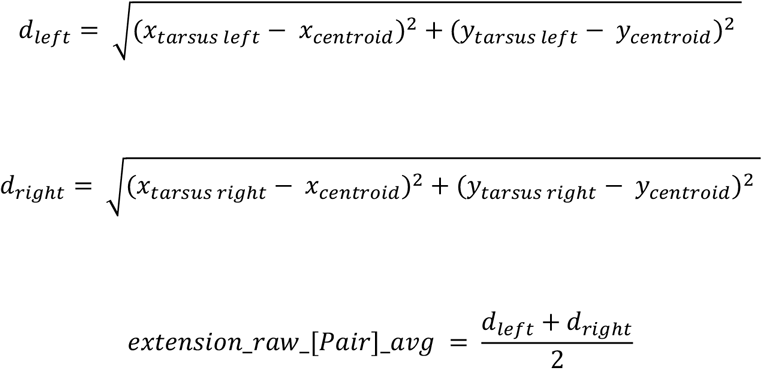

For Floor-oriented Standing frames, we also calculated left-right asymmetry magnitudes for the eyes, thorax, abdomen, wings, and each leg pair using the precomputed x- and y-asymmetry indices, and summarized these as an overall left-right asymmetry score. For Standing frames with Wall-Side Orientation prediction ≥ 0.75, top-bottom asymmetry magnitudes for the head, thorax, and abdomen were calculated and summarized as overall top-bottom asymmetry scores.

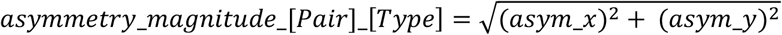

Two overall postural asymmetry features (overall_asymmetry_LR and overall_asymmetry_TB) were calculated as the mean of all LR or TB asymmetry magnitudes.

Wing-spread and left-right leg-pair spread angles were summarized within selected Standing, Walking, and Groom-Wings contexts by calculating the mean and 95th percentile of each angle feature across valid frames.

Kinematic features were calculated from keypoint-derived instantaneous movement signals within valid BxO contexts. Most Behaviors were separated into Floor, Wall-Side, and Ceiling Orientation contexts, while Jump and Scramble retained as collapsed orientation-independent contexts. Wall-Up and Wall-Down were excluded from kinematic features. Instantaneous kinematic signals included body translation, acceleration, and jerk; angular velocity, acceleration, and jerk; walking curvature; head movement; and front-, middle-, and back-leg translational and angular movement. Head movement was derived from five head keypoints by dropping the lowest-likelihood head keypoint on each frame and averaging the remaining signals.

Kinematic features were summarized as bout-based features by first summarizing each bout individually and then aggregating those summaries across bouts of the same type within each animal. Non-walking bouts were summarized over the full bout, whereas walking bouts were summarized using a trimmed core window together with separate start and end windows of five frames each. For each bout, we calculated the mean, standard deviation, median, Pearson’s non-parametric skewness, and robust minimum and maximum values defined as the mean of the bottom and top 5% of values. Only features derived from at least three valid bouts were retained. We also calculated a global locomotor summary feature by averaging whole-body speed across all frames for each animal.

To calculate transition probabilities for Behavior labels and combined BxO labels, we identified successive transitions in which one label was followed by a different label, such that each transition had a “From” and “To” state. If *N*_*From*>T*o*_ is the number of transitions from the From state to the To state, and *N*_*Total From*_ is the total number of outgoing transitions from that From state, then transition probability was calculated as:

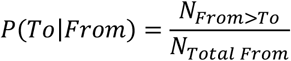

When calculating transitions for the BxO combined labels, we collapsed all Jump labels into a single category due to their rarity, and collapsed all Scramble labels into a single category due to the rapid transitions between Orientations. Transition features supported by fewer than six outgoing transitions from the source state were masked downstream.

Spatial and directional features were calculated from the centroid position and body orientation of the animal. First, we converted the centroid to a rotationally-invariant polar coordinate system, where (299.5, 299.5) is considered the chamber center:

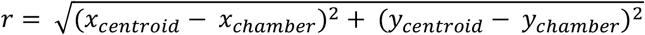

We then used the chamber radius to calculate a normalized radius *r*_*norm*_, where the center is 0 and the wall is 1:

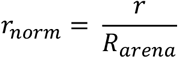

For each behavior or BxO combination, we collected all *r*_*norm*_ values and calculated radial-position statistics including mean, median and standard deviation.

We calculated relative angle, where 0° points directly away from the chamber center and 180° is pointing directly towards the chamber center, by subtracting the positional angle *θ*_*po*s_ from its actual body angle *α*_*fly*_:

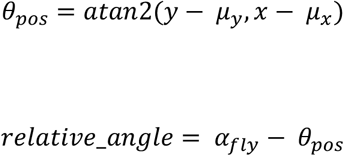

We calculated center and periphery occupancy fractions by classifying normalized radial positions as center (*r*_norm_ ≤ 0.4) or periphery (r_norm_ ≥ 0.7) and computing the proportion of frames in each region within selected behavioral contexts.

We also quantified directional orientation relative to the chamber center by calculating the framewise angle between the body axis and the radial position vector. After excluding frames near the chamber center (*r*_norm_ < 0.1), we summarized the mean and standard deviation of the absolute relative angle within each behavioral context.

